# Transcriptional reactivation of the tRNA^Ser^/tRNA^Tyr^ gene cluster in *Arabidopsis thaliana* root tip (V2)

**DOI:** 10.1101/2023.09.27.559738

**Authors:** Guillaume Hummel, Priyanka Kumari, Chenlei Hua, Long Wang, Yan-Xia Mai, Nan Wang, Negjmedin Shala, Emir Can Kaya, Jean Molinier, Jia-Wei Wang, Chang Liu

**Author notes:** These authors contributed equally to this work (Guillaume Hummel, Priyanka Kumari). Corresponding contacts: Hummel, Guillaume, Liu, Chang. The authors responsible for the distribution of materials integral to the findings presented in this article, and in accordance with the policy described in the Instructions for Authors, (https://academic.oup.com/plphys/pages/General-Instructions) are: Guillaume Hummel and Chang Liu.

## Abstract

Plants retain a repetitious tRNA gene content in their nuclear genome. How important are these individuals, how exactly plants orchestrate their usage, and for what purposes, is poorly understood. *Arabidopsis thaliana* chromosome 1 holds a cluster of tandemly repeated serine– and tyrosine-decoding tRNA genes (SYY cluster). They intersect with constitutive heterochromatin and are silenced in most parts of the plant. Yet, the natural conditions leading to their transcription remain unknown. Here, we resolve the tissular expression pattern of this cluster along seedling establishment. We show that the root cap columella and few adjacent lateral root cap cells are the main sources of SYY cluster tRNAs. The transcriptional reactivation of the SYY cluster occurs in these tissues although elevated DNA methylation levels. Furthermore, we evidence that these cells are able to accumulate high levels of a transgenic glycoprotein rich in serine, tyrosine, and proline, and that the CRISPR/Cas9 deletion of the SYY cluster alters the phenomenon. Altogether, our work sheds light on pioneering evidence of a developmental and cell-specific expression program for a plant tRNA gene. We provide new perspectives on the role of peculiar tRNA genes in conferring a potential for the high synthesis of glycoproteins in protective tissues of the meristem.

## INTRODUCTION

Transfer RNAs (tRNAs) represent a category of small non-coding RNA species. Found in all life forms, tRNAs are evolutionarily ancient RNAs (Chan and Lowe, 2016). Their existence relates to an essential process for life: protein biosynthesis. tRNAs work as adaptor molecules that charge an amino acid at their 3’ ends, and subsequently transfer it at the C-terminus of nascent polypeptides inside ribosomes (Hoagland et al., 1958; Berg et al., 1961; Holley et al., 1965). The fidelity of translation relies on highly conserved structural elements (Berg and Brandl, 2021; Giegé and Eriani, 2023). They ensure that tRNAs are charged with the cognate amino acid, and accurately match it with one of the 61 codons composing messenger RNAs (mRNAs) (Nirenberg et al., 1965). Therefore, a set of tRNA species exists for the decoding of each amino acid (Chan and Lowe, 2016).

The evolution of nuclear genomes went along with a complexification of tRNA gene (tDNA) contents, in terms of copy number and sequence diversity (Goodenbour and Pan, 2006; Santos and Del-Bem, 2022). tRNAs are generally represented by multigenic tDNA families from which the individual members are distributed in diverse chromosomal locations (Hummel and Liu, 2022; Santos and Del-Bem, 2022). Numerous mutational events accompanied the genomic proliferation of tDNAs and diversely increased the pool of unique tRNA sequences among species (Goodenbour and Pan, 2006). Polymorphisms either do not affect the predicted structure of tRNA transcripts, or make them variably diverging from a functional tRNA (Chan and Lowe, 2016). Named “tRNA-like”, these pseudo-RNAs might be impaired in expression, stability and/or their function in translation.

The copy number of a nuclear tDNAome appears positively correlated to the genome size, the content of protein-coding genes and codon frequencies (Goodenbour and Pan, 2006; Michaud et al., 2011; Santos and Del-Bem, 2022). It implies that nuclear genomes retain multiple copies of corresponding tDNAs in proportion to the presumed demand for specific amino acids, ensuring an ample supply of tRNAs to meet the requirements of protein synthesis. Nonetheless, a growing body of evidence reveals that in a given species and condition, only a restricted fraction of the nuclear tDNAome might be transcribed. Immunoprecipitations of chromatin bound by the RNA polymerase (RNAP) III machinery, and tRNA sequencing experiments, showed that roughly half of the 619 tDNAs contribute to less than 1% of the tRNA abundance in human (Canella et al., 2010; Moqtaderi et al., 2010; Oler et al., 2010; Gogakos et al., 2017; Thornlow et al., 2018; Torres, 2019). Similar conclusions were recently stated in plants (Hummel et al., 2020; Liu and Sun, 2021; Ma et al., 2021; Hummel and Liu, 2022). Therefore, it is questionable why eukaryotes retain so much tDNAs in their nuclear genome. Are some of them developmentally/environmentally regulated? What is the biological importance of individual tDNAs?

The nuclear genome of the plant *Arabidopsis thaliana* Col-0 consists of five chromosomes carrying 586 tDNAs (Cognat et al., 2022). While most are dispersed along arms, a subset of proline (Pro), serine (Ser) and tyrosine (Tyr) tDNAs organize in clusters (Green and Weil, 1989; Beier et al., 1991; Theologis et al., 2000; Michaud et al., 2011; Hummel et al., 2020; Hummel and Liu, 2022). Chromosome 1 (Chr1) and Chr2 comprise six small clusters (*c.a.* 3-6kbps) numbering up to 11 tDNAs^Pro^ (Hummel et al., 2020; Hummel and Liu, 2022). Furthermore, Chr1 holds a large array (*c.a.* 40kbps) of 27 tDNAs^Ser^ and 54 tDNAs^Tyr^ tandemly repeated 27 times (Ser-Tyr1-Tyr2)_27_ (Hummel et al., 2020; Hummel and Liu, 2022), which is referred in this work as the “SYY cluster”. A previous study revealed that tDNA clusters stand in epigenomic environments unfavourable for RNAPIII transcription and are silent (Hummel et al., 2020). There is currently no clue as to what triggers their expression and to what biological end. Herein, we explore the conditions, molecular mechanisms, and physiological significance of their reactivation by focusing on the SYY cluster.

## RESULTS

### The SYY cluster consists of a core region which expanded by tandem duplication

Out of the 20 tDNA families encountered in the Col-0 nuclear genome, tDNA^Tyr^ is the most abundant, and counts 70 genes (Cognat et al., 2022). For the sake of description, and to explore their genetic diversity and organization, we sought to hierarchize tDNAs^Tyr^ based on three factors: the layout (dispersed: D/clustered: C), nucleotide polymorphisms (NPs), and gene copy numbers (**Figs. 1A and S1A**). Inspired from 5S ribosomal DNA (rDNA) work (Cloix et al., 2002), the nomenclature defines tDNA NPs represented by the highest gene copy number as “major”. All others are decreasingly categorized as “minor”. Thus, D-tDNAs^Tyr^ comprise a MAJOR represented by 14 copies and two MINORs represented by a total of two copies (**Fig. 1A**). C-tDNAs^Tyr^ comprise a MAJOR represented by 32 copies and nine MINORs represented by a total of 22 copies (**Fig. 1A**).

**Figure 1.**
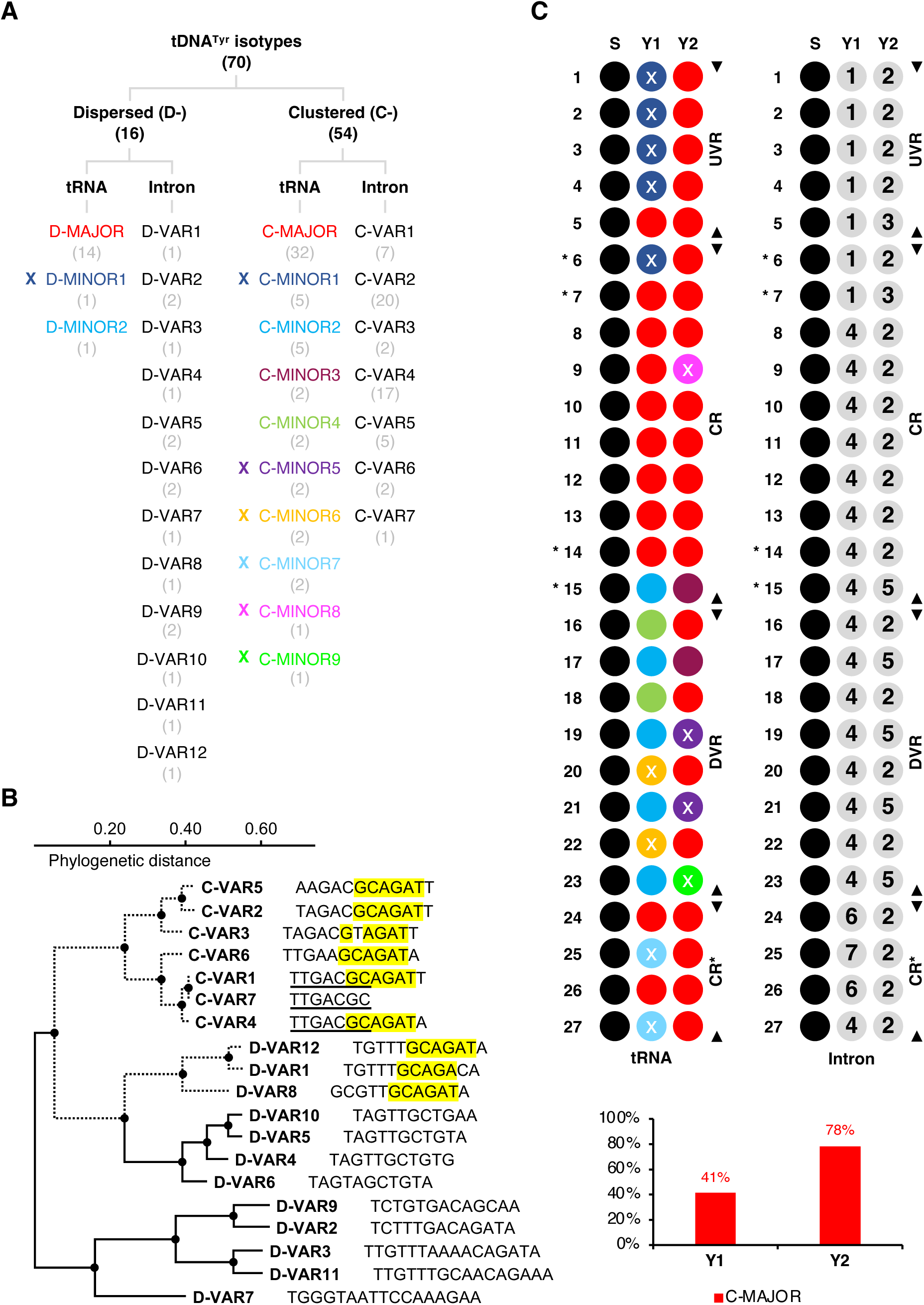
Diversity and organization of tRNA^Tyr^ genes in the Col-0 nuclear genome. **A**, Nuclear tDNA^Tyr^ hierarchy tree. Numbers in brackets refer to the gene copy number of each sub-category. Crosses flag the sub-categories represented by tRNA-like pseudogenes. **B**, Nuclear tDNA^Tyr^ intron phylogeny. Dotted branches show the common ancestry between the two clades comprising D-VARs 1/8/12 and C-VARs. Corresponding sequences are characterized by a conserved GCAGAT motif highlighted in yellow. CVAR7 results from the truncation of an ancestral intron commonly shared with C-VARs 1 and 4. The conserved sequence is underlined. **C**, Distribution of tDNAs^Tyr^, and their introns, within the Col-0 SYY cluster. Colour and numerical legends are identical to A. The location of tRNA-like pseudogenes is indicated by crosses. The core region (CR), upstream and downstream variable regions (UVR and DVR respectively), and the CR* are delimited with black arrows. The SYY pairs associated with the tandem expansion of the CR (*i.e.,* 6, 7, 14 and 15) are flagged with an asterisk. Histograms present the proportion of genes bearing the C-MAJOR tRNA^Tyr^ sequence at Y1 and Y2 positions.

The analysis of secondary structures revealed that out of the 16 D-tDNAs^Tyr^, 15 lead to functional tRNAs and only one (*i.e.,* MINOR1) leads to a tRNA-like transcript (**Figs. 1A and S1B**). By opposition, out of the 54 C-tDNAs^Tyr^, a total of 13 MINORs (*i.e.,* 1, 5, 6, 7, 8 and 9) lead to tRNA-like RNAs (**Fig. 1A and S1C**). These pseudo-tRNAs are characterized by different structural mismatches which might be deleterious for their stability and role in translation (**Fig. S1B and C**). They are probably non-functional, and might be conveyed to degradation by tRNA quality controls (Phizicky and Hopper, 2023).

tDNAs^Met^ and tDNAs^Tyr^ are the only tDNA species containing an intron in the nuclear genome of *A. thaliana* (Michaud et al., 2011). We set up a secondary nomenclature based on the sequence of these introns (**Fig. 1A and B**). D-tDNAs^Tyr^ have 12 different intron variants (VARs), while C-tDNAs^Tyr^ have only seven. Interestingly, we found a highly conserved GCAGAT motif in C-VARs (**Fig. 1B**). All C-tDNAs^Tyr^ have the motif except one which underwent a truncation (*i.e.,* C-VAR7, **Fig. 1A and B**). The motif is also present in D-VARs 8 and 12 species, which are represented by one C-tDNA^Tyr^ each (**Fig. 1A and B**). This suggests that during the evolution of tDNA^Tyr^ content in Brassicaceae genomes, the SYY cluster is phylogenetically linked to these two D-tDNA^Tyr^ species and originated from the tandem duplication of one or more common tDNA ancestors.

Then, we confronted the two nomenclatures to unveil the internal organization of the SYY cluster (**Fig. 1A and C**). We assigned to each repeat unit (1 to 27) and to each tDNA^Tyr^ cassette (Y1 and Y2) the associated C-MAJOR/C-MINOR and intron C-VAR (**Fig. 1C**). The resulting pattern led us to the identification of an obvious region, where the pairs of C-MAJOR/C-MAJOR concentrate. We defined it as the core region (CR). This latter is flanked by variable regions in which the proportion of C-MINORs and tDNA-like pseudogenes increases. Besides, at one side of this cluster region, several copies of the C-MAJOR/C-MAJOR pairs are present, which we named CR*.

As a whole, the C-MAJOR is mainly represented by the Y2 position (*c.a.* 80%) while CMINORs are preponderantly found at the Y1 cassette (*c.a.* 60%) (**Fig. 1C**). Due to the presence of unique sequence variations, the transcripts derived from D– and C-MAJORs can be specifically probed, allowing to study the expression and regulation of C-tDNAs^Tyr^ in this cluster (Hummel et al., 2020).

### Root tip cells source SYY cluster transcripts during vegetative development

It has been previously shown that the SYY cluster is embedded in constitutive heterochromatin (CH) and thus, epigenetically silenced (Hummel et al., 2020). It remains to be deciphered whether particular growth conditions would allow the transcription of this SYY cluster.

Developmental transitions have been pivotal in the understanding of CH dynamics (Benoit et al., 2013; Simon and Probst, 2023). Germination associates with a decompaction of CH promoting the expression of the silent fraction of 5S rDNA clusters in *A. thaliana* cotyledons (Mathieu et al., 2003). Thus, we asked as an outset whether CH reshaping during germination releases the silencing of the SYY cluster. We monitored its expression in six timepoints corresponding to imbibed seeds (0 days after stratification, DAS), testa rupture (1DAS), endosperm rupture and radicle emergence (2DAS), cotyledon emergence and greening (3DAS), established plantlets (4DAS) and adult plants (14DAS) (**Fig. 2A**).

**Figure 2.**
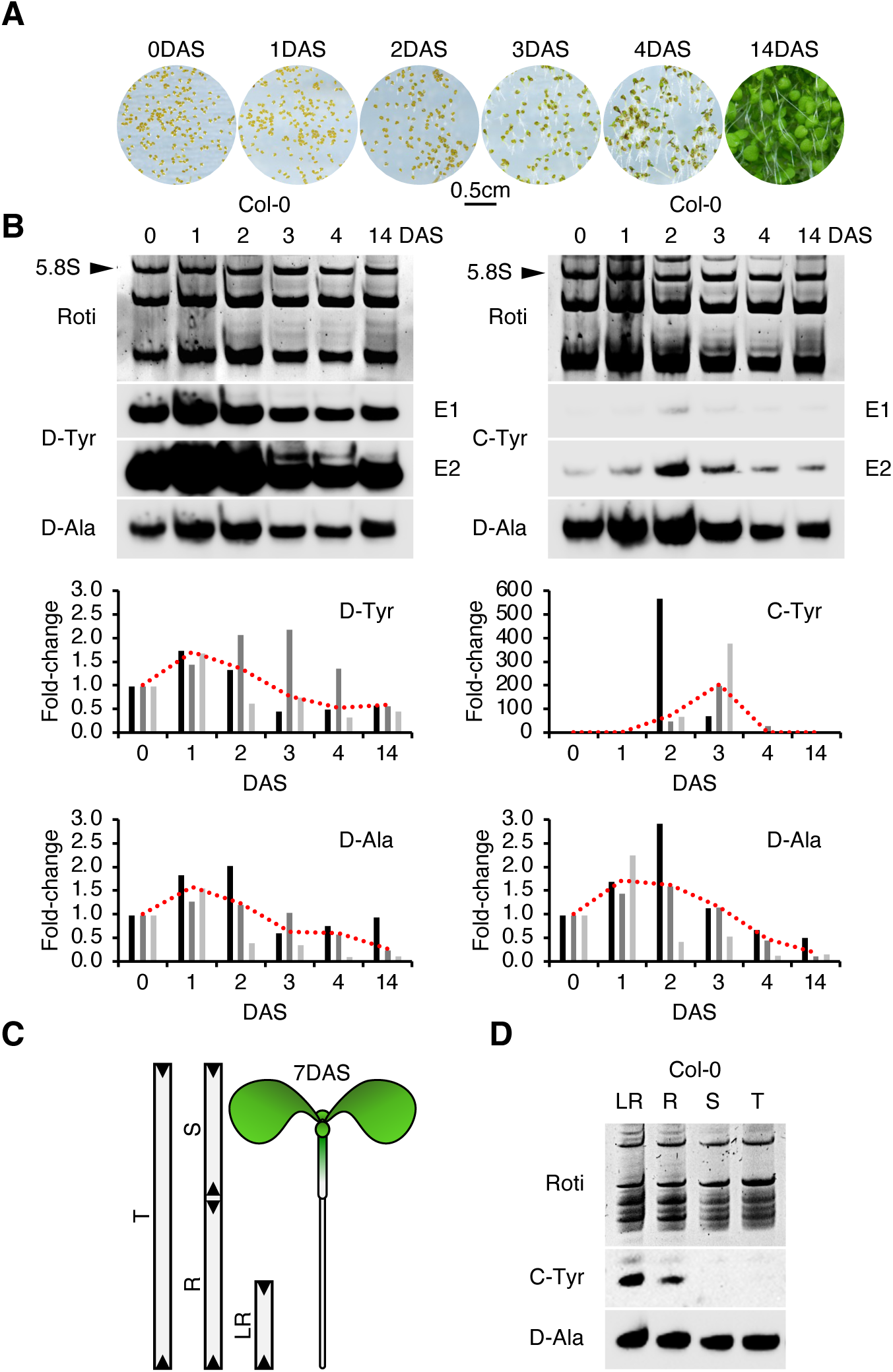
Expression pattern of the SYY cluster during vegetative development. **A**, Representative thumbnails of 0, 1, 2, 3, 4, and 14DAS germination timepoints. **B**, Northern blot detection of D-tRNAs^Tyr^ (D-Tyr probe), C-tRNAs^Tyr^ (C-Tyr probe), and D-tRNAs^Ala^ (D-Ala probe) in the total RNA extracts of consecutive germination timepoints. Two equal exposures (E1, 2) are presented for D/C-tRNAs^Tyr^. A representative of three independent runs is every time depicted. Graphs present the quantification analysis of D-Tyr, C-Tyr, and D-Ala probe signals in three independent biological replicates. After a calibration based on the 5.8S rRNA staining (see arrows), results were expressed in fold-change using the 0DAS timepoint as a reference (see histograms). Red dotted lines connect the median value of each timepoint. **C**, Segmentation of a 7DAS plantlet. T: total, S: shoot, R: root, LR: lower root. **D**, Northern blot detection of C-tRNAs^Tyr^ and D-tRNAs^Ala^ in the total RNA extracts of different organs.

Expectedly, D-tRNAs^Tyr^ accumulate during germination and remain dominant among tRNA^Tyr^ species (**Fig. 2B**). C-tRNAs^Tyr^ stay lowly detectable but their steady-state level increases transiently in 2DAS and 3DAS plantlets (**Fig. 2B**). As an additional control, membranes were stripped and rehybridized with a D-tRNA^Ala^ probe. D/C-tRNAs^Tyr^ and DtRNAs^Ala^ were quantified from unsaturated exposures with ImageJ, and signals normalized based on the RNAPI transcription-derived 5.8S rRNA quantified from Roti staining. 0DAS signals were used as reference to calculate fold-changes. The results indicated that C-tRNAs^Tyr^ momentarily accumulate in 2/3DAS plantlets. Fold-changes vary between 54 and 572 for the 2DAS stage and between 76 and 382 for 3DAS stage (**Fig. 2B**). Only for seed batch #2, C-tRNAs^Tyr^ were detected in 4DAS plantlets but fold-change (34) remained lower than abovementioned (**Fig. 2B**). By contrast, the accumulation of D-tRNA^Tyr^ and DtRNA^Ala^ species is similar, and characterized by a slight increase at 1 and 2DAS followed by a progressive decrease till 14DAS. Fold-changes only varying between 0.35 and 2.21 for D-tRNAs^Tyr^ and between 0.13 and 2.94 for D-tRNAs^Ala^, the steady-state levels of these two species are relatively stable compared to C-tRNAs^Tyr^. Thus, C-tRNAs^Tyr^ show a temporal accumulation pattern restricted to 2/3DAS plantlets. Nonetheless, the upregulation of the SYY cluster remains very low compared to D-tDNAs^Tyr^ expression (**Fig. 2B**).

To infer whether the transient detection of C-tRNAs^Tyr^ during germination comes from the enrichment of RNAs from specific source tissues in 2/3DAS extracts, we further analysed RNA extracted from different tissues, which included complete seedlings (T), shoots (S), whole roots (R), and the lower half of roots (LR). While control D-tRNAs^Ala^ accumulate equally among the four samples, we noted a preferential enrichment of C-tRNAs^Tyr^ in R and LR RNA extracts (**Fig. 2D**). Nevertheless, they are nondetectable in T and S RNA extracts. Remarkably, their enrichment is higher in the lower part of the root comparatively to the whole root organ (**Fig. 2D**). It suggests that a gradient pattern governs the SYY cluster expression along the root axis. Altogether, these results disclose that at least one tissue located in the lower part of roots abundantly accumulate C-tRNAs^Tyr^ during the vegetative development of *A. thaliana*.

### The SYY cluster is indeed the genomic region producing C-tRNAs^Tyr^ in roots

To ascertain that the SYY cluster is the genomic region responsible for the accumulation pattern of C-tRNAs^Tyr^ in roots, knockout mutants bearing its deletion were produced using the CRISPR/Cas9 technology (**Fig. 3**). Up(region U) and downstream (region D) of the SYY cluster were used as guide RNA (gRNA) (**Fig. 3A**). We parallelly used two approaches relying on either dual or multiplexed guide RNA (gRNA) layouts (**Fig. 3B**) (Zhang et al., 2016; Wu et al., 2018). In the first, one gRNA was designed by each flanking region (GH1 to 2, **Fig. 3B**). In the second, three gRNAs were designed by region (LW1 to 6, **Fig. 3B**). The outcome of repair of the CRISPR/Cas9 SYY cluster targeting was screened by PCR using two primer sets. The couple 1+2 only amplifies if the SYY cluster is present (**Fig. 3A and C**). Noteworthily, oligonucleotide 1 hybridizes the truncated C-VAR7 intron uniquely represented in the Col-0 genome (**Figs. 1 and 3A**). The couple 3+2 only works if the SYY cluster is deleted (**Fig. 3A and C**). We systematically obtained PCR products shorter than what putatively expected with the 1+2 couple in WT (**Fig. 3C**). In addition, internal primers α to ζ allow to screen for the presence/absence of the three C-tDNA cassettes (S, Y1, and Y2, **Fig. 3A and D**).

**Figure 3.**
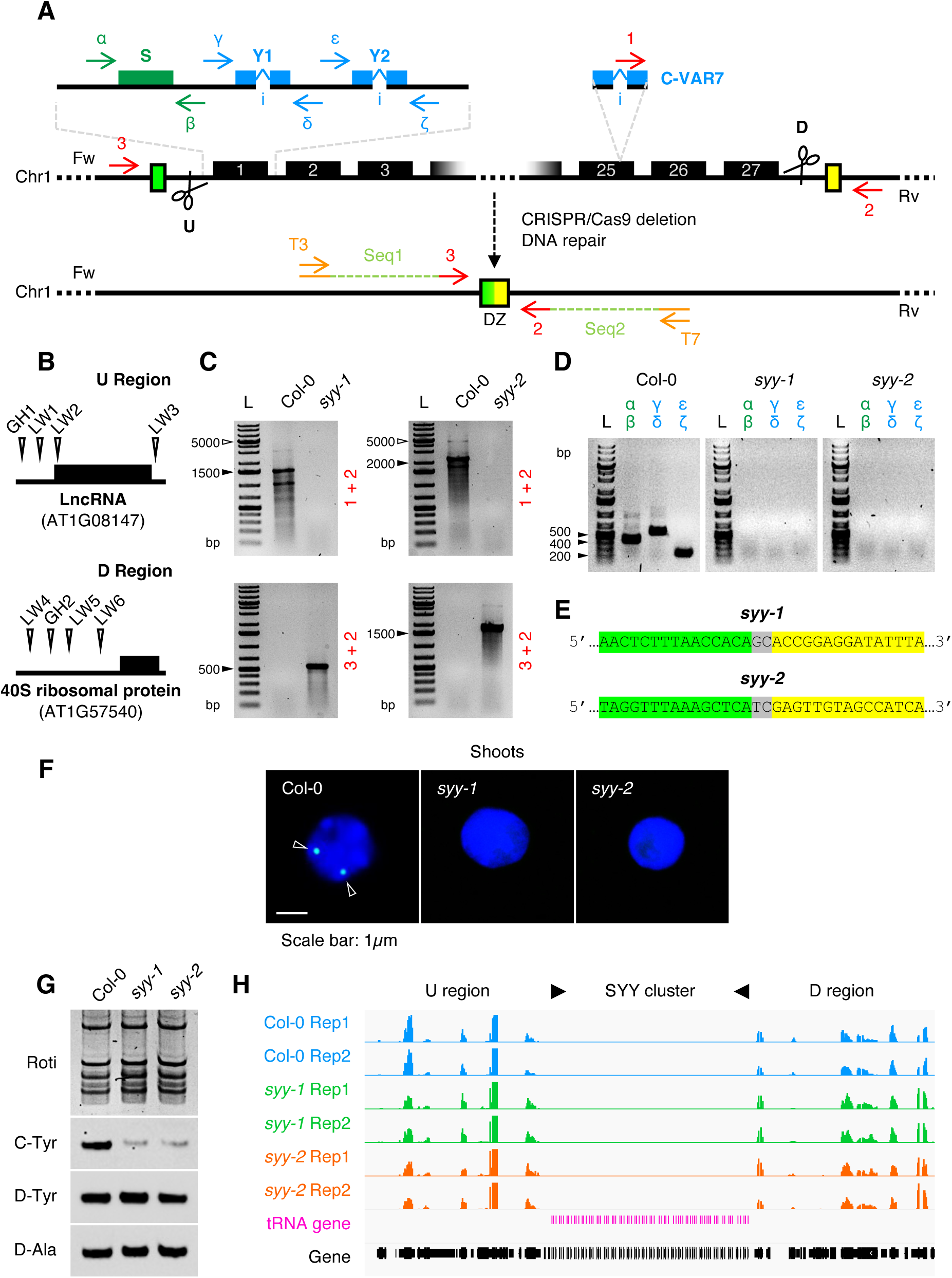
Generation and characterization of tRNA gene deletion alleles. **A**, Screening procedure for lines lacking the SYY cluster. Dual (“GH”) and multiplexed (“LW”) gRNAs used in U/D regions are detailed in **B**, with proximal genomic entities. DZ, deletion zone. **C and D**, PCR profile of homozygous lines. Black arrows point ladder (L) sizes the closest from amplicons, while white ones indicate the closest sizes of putative amplicons according to TAIR10 genome. 1+2 = 4543bps (*syy-1* screen) or 5228bps (*syy-2* screen). 3+2 = 547bps (*syy-1* screen) or 1875bps (*syy-2* screen). α+β = 408bps. γ+δ = 564bps. ε+ζ = 212bps. **E**, Sequence of *syy-1* and *syy-2* alleles. Few nucleotides up and downstream (green and yellow regions in A) of the DNA junction (GC/TC di-nucleotides) are shown. **F**, Subnuclear detection of the SYY cluster in Col-0, *syy-1* and *syy-2* shoot 2C nuclei. BAC-FISH signals are identified with arrows. Pictures are representatives of around 100 observations. **G**, D/C-tRNAs^Tyr^ and D-tRNAs^Ala^ steady-state levels in Col-0, *syy-1* and *syy-2* 7DAS roots. **H**, Expression level of genes flanking SYY cluster deletion (see U and D regions in A).

The deletion zone (DZ) of positive plants was amplified with tailed 3 and 2 primers holding a trigger (T3/7 respectively) and trailer sequence (seq1/2 respectively) (**Fig. 3A**). The sequencing of amplicons led to the identification of two alleles we called *syy-1* and *syy-2* (**Figs. 3E and S2**). *syy-1* originates from the duplexed gRNA approach, excludes the upstream long non-coding RNA gene, but keeps the downstream 40S ribosomal protein gene (**Fig. S2**). *syy-2* originates from the multiplexed gRNA approach, and preserves the two flanking loci (**Fig. S2**). Mapping DZ up– and downstream sequences (green and yellow respectively, **Fig. 3A**) to the TAIR10 reference showed that repair occurred at GC (*syy-1*) or TC (*syy-2*) dinucleotides common to both side extremities (**Figs. 3E and S2**).

We further characterized SYY cluster deletion at the cytogenetical level, using fluorescent *in situ* hybridization (FISH). The hybridization and detection of a digoxigenin-labelled BAC covering the SYY cluster (*i.e.,* F9K23) led to two FISH signals in 2C nuclei from wild-type plants (**Fig. 3F**). On the contrary, no FISH signals were detected in *syy-1* or *syy-2* nuclei, confirming the absence of the SYY cluster in these lines.

At the RNA level, C-tRNAs^Tyr^ accumulate in Col-0 roots but deplete in *syy-1* and *syy-2* ones, whereas D-tRNAs^Tyr^ and control D-tRNAs^Ala^ accumulate comparatively among the three genotypes (**Fig. 3G**). A faint signal persists in mutant lines, what we interpret as a background of cross-hybridization. Importantly, we verified by poly-A RNA sequencing that SYY cluster deletion did not alter the local gene expression pattern in U and D regions (**Fig. 3H**).

These experiments demonstrate that the SYY cluster is the sole region producing C-tRNAs^Tyr^ in our seed’s genome, and responsible for the developmental expression pattern described in **Fig. 2**.

### Columella and border-like cells are the main expression sites of the SYY cluster

To identify the tissues expressing the SYY cluster in LR sections (**Fig. 2D**), we performed whole-mount *in situ* hybridization (WISH) coupled with tyramide signal amplification (TSA) to study the distribution of C-tRNAs^Tyr^ transcripts in 4DAS root tissues (Andras et al., 2001; Wójcik et al., 2018).

The WISH experiments generated a staining pattern consistent with the results obtained by northern blot (**Fig. 2D**), in which the signal intensity progressively increased along roots (**Fig. 4A**). The counter-lighting made evident that the staining is homogenous in the division zone and the tip. Thus, the tissular identity is not the main biological parameter for the establishment of this gradient but more the fact that cells are dividing and not fully differentiated. Nevertheless, the four peripheral cells of the columella (RCC1-4), and few adjacent border-like cells (LRC), were systematically subject to a staining higher than dividing cells (**Fig. 4B**) (Dolan et al., 1993). As expected, control *syy-1* rootlets didn’t exhibit the staining (**Fig. 4A and B**).

**Figure 4.**
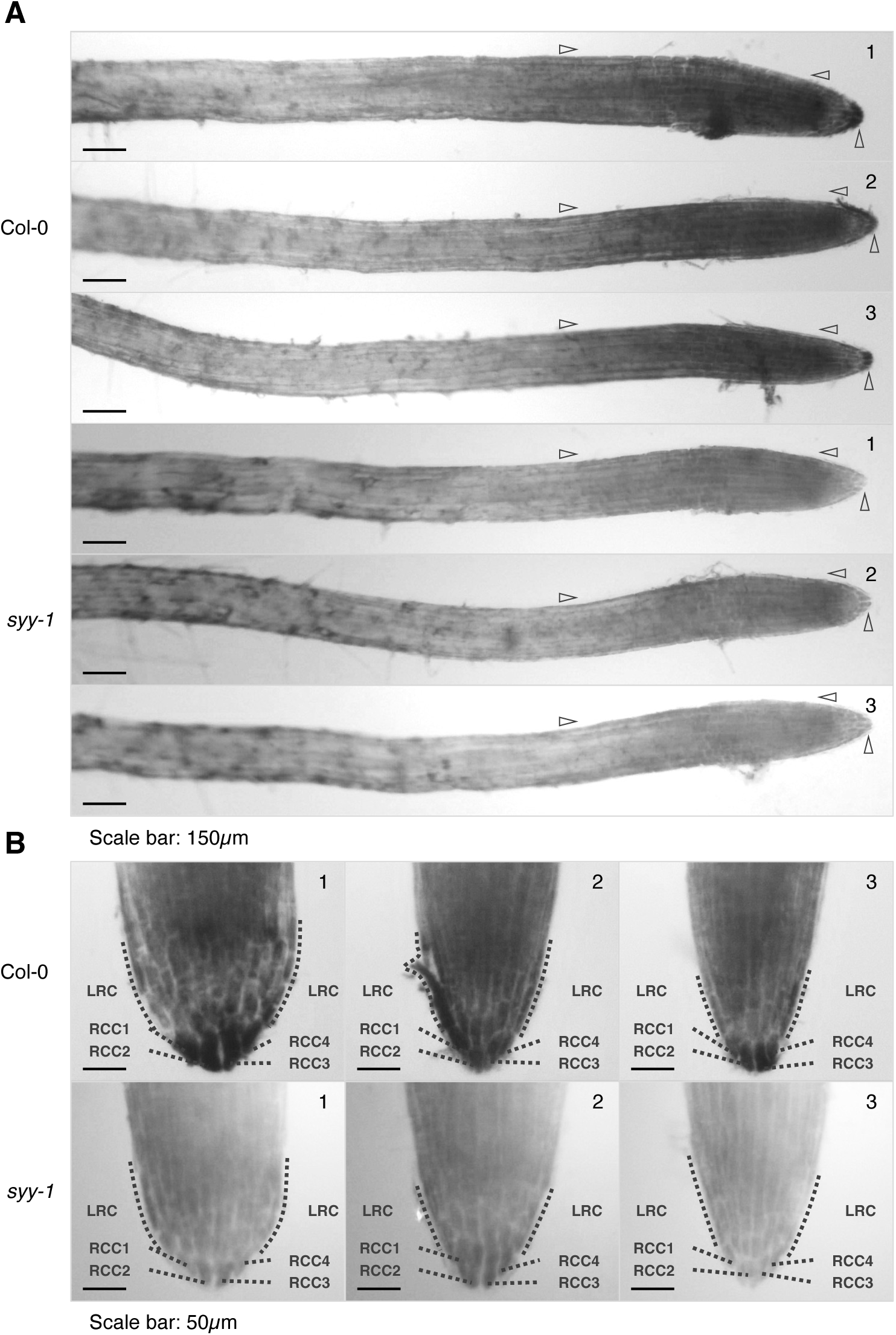
Expression pattern of the SYY cluster in root tissues. **A**, WISH-TSA detection of C-tRNAs^Tyr^ in 4DAS Col-0 and *syy-1* rootlets. Arrows delimitate zones of differential staining between genotypes. **B**, Enlargements of 4DAS Col-0 and *syy-1* tips. The four external columella (RCC1-4) and lateral root cap (LRC) cells with a strong C-tRNAs^Tyr^ accumulation are titled with lines. Three representatives are depicted.

### C-tDNAs^Tyr^ are functional and constitutively expressed in roots upon transgenesis

The functionality of C-tDNA^Tyr^ gene structure was only deduced from the analysis of their *cis*-motifs, but lacked an experimental validation *in vivo* (Hummel et al., 2020). We took advantage of CRISPR/Cas9 lines lacking the SYY cluster to reintroduce back transgenic C-tDNAs^Tyr^ and study their transcription.

We either cloned the first repeat unit in upstream of the SYY cluster (1xSYY), or engineered a C-tDNA^Tyr^ (Ing-Y) (**Fig. 5A**). The latter contains the MAJOR sequence and the C-VAR2 intron (most represented in the SYY cluster), combined to the promoter and terminator of a D-tDNA^Tyr^ (AT5G61835) bearing all *cis*-elements required for an optimal transcription of plant nuclear tDNAs (**Figs. 1A and 5B**) (Yukawa et al., 2000; Michaud et al., 2011). Constructs were stably agrotransformed in the *syy-1* background. Three *syy-1* Ing-Y and two *syy-1* 1xSYY complemented lines were selected. The proportion of fluorescent seeds in T2 is classic for the segregation of a single transgenic copy, except for *syy-1* Ing-Y #1 and *syy-1* 1xSYY #2 displaying a slight distortion (**Fig. 5C**). Nevertheless, the Southern blotting of Ing-Y and 1xSYY transgenes made clear that all these lines are multi-insertional (**Fig. 5D**).

**Figure 5.**
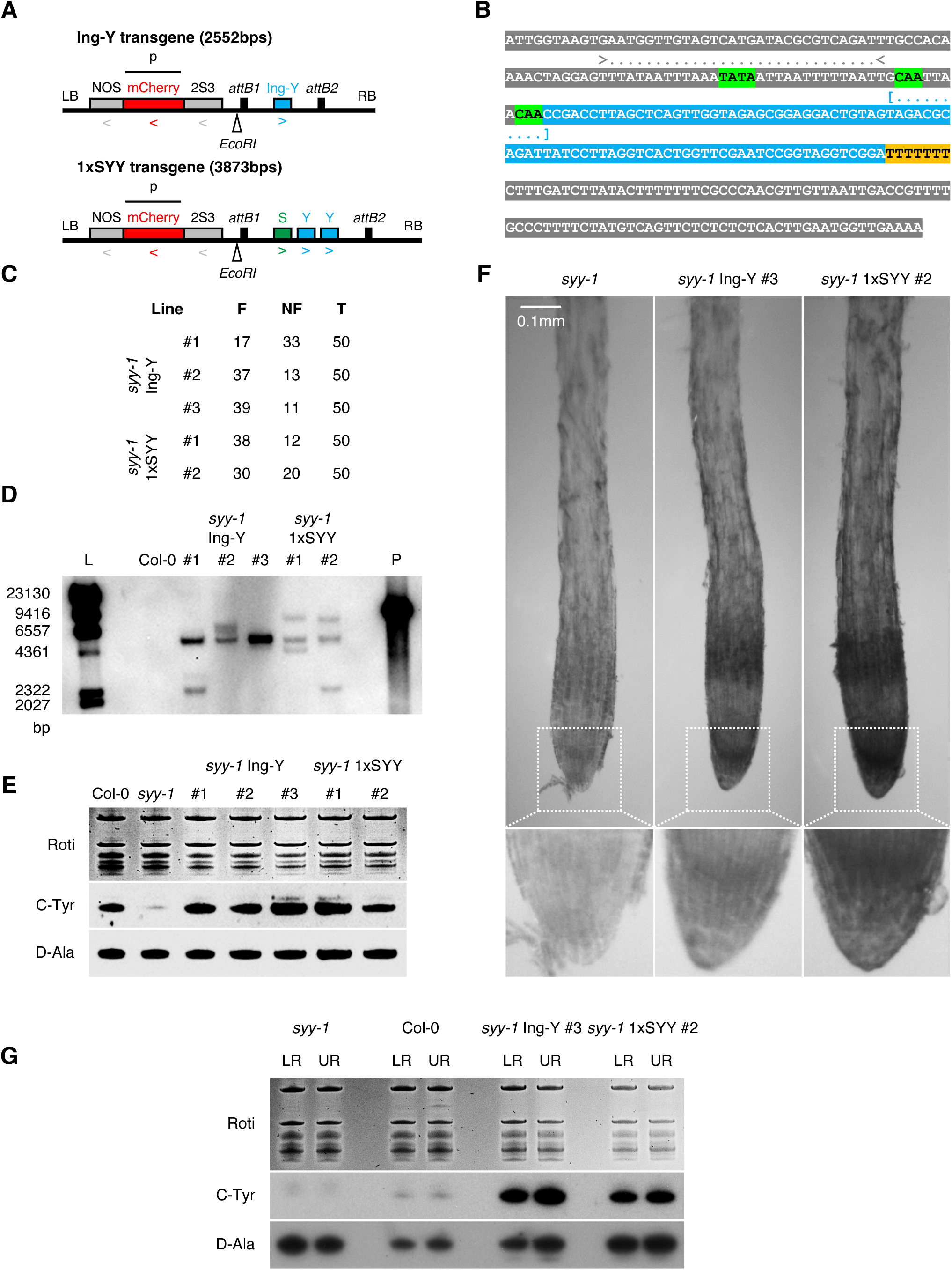
Stable agro-transformation of transgenic tRNA genes. **A**, Genetic map of constructs used for *syy-1* complementation, together with the transcriptional orientation (arrows), restriction enzyme site (*EcoRI*) and probe (p) used for Southern blotting. LB: left border, RB: right border, Ing-Y: engineered C-tDNA^Tyr^, 2S3: 2S albumin gene 3 promoter, NOS: nopaline synthase terminator. **B**, Sequence of the Ing-Y gene. The blue sequence corresponds to the major C-tRNA^Tyr^. Brackets delimitate the intron. Within upstream region, the A/T-rich region is demarked with arrows. Regulatory TATA-like, CAA, and poly-T *cis*-motifs are highlighted in green and orange respectively. **C**, Segregation of non-fluorescent (NF) and fluorescent (F) seeds in *syy-1* Ing-Y and *syy-1* 1xSYY lines. **D**, Southern blotting of Ing-Y and 1xSYY transgenes in the digested genomic DNA of Col-0 (negative control) and complemented *syy-1* lines. The undigested plasmid (P) used to amplify the p probe was used as positive control. L: ladder. **E**, C-tRNAs^Tyr^ and D-tRNAs^Ala^ steady-state levels in Col-0, *syy-1*, *syy-1* Ing-Y (three independent lines), and *syy-1* 1xSYY (two independent lines) 7DAS roots. **F**, WISH detection of C-tRNAs^Tyr^ in 4DAS *syy-1*, *syy-1* Ing-Y and *syy-1* 1xSYY rootlets. **G**, Northern blot detection of C-tRNAs-^Tyr^ and D-tRNAs^Ala^ in the lower (LR) and upper (UR) root extracts of 7DAS *syy-1*, Col-0, *syy-1* Ing-Y, and *syy-1* 1xSYY plants.

For both types of transgenic lines, the accumulation of C-tRNAs^Tyr^ is restored in total root RNA extracts compared to *syy-1* (**Fig. 5E**). By contrast, the steady-state level of control D-tRNAs^Ala^ does not vary (**Fig. 5E**). The introduction of SYY expression cassettes resulted in much higher C-tRNAs^Tyr^ expression than the level offered by the endogenous SYY tandem repeat with 34 copies. This is due to their constitutive expression along root, as showed by WISH with *syy-1* 1xSYY #2 and *syy-1* Ing-Y #3 lines (**Fig. 5F**). Interestingly, no particular enrichment of the C-tRNA^Tyr^ was observed in the LRC and RCC (**Figs. 4 and 5F**). It further confirms that the natural SYY cluster is subject to a developmental regulation permitting the upregulation of corresponding loci in these tissues. The constitutive expression of transgenic tDNAs in different parts of roots was further verified by northern blotting (**Fig. 5G**). C-tRNAs^Tyr^ are strongly detected in lower and upper root RNA extracts of *syy-1* Ing-Y #3 and *syy-1* 1xSYY #2 lines, while a faint and no signal are observed in Col-0 and syy-1 ones (**Fig. 5G**). By contrast, control D-tRNAs^Ala^ accumulate comparatively among all genotypes (**Fig. 5G**).

These results indicate that despite being mainly silenced *in vivo*, C-tDNAs^Tyr^ are functional gene loci, able to be strongly and constitutively expressed.

### High DNA methylation levels accompany natural SYY expression in roots

Knowing that the SYY cluster reactivates along Col-0 roots, we sought to decipher how it relates to changes in its methylation landscape. Indeed, DNA methylation is an epigenetic mark involved in SYY cluster silencing (Hummel et al., 2020). We performed chop-PCR assays with methylation-sensitive restriction enzymes (MSREs) and bisulfite sequencing data re-analysis to this end.

The alignment of C-tDNA^Tyr^ sequences led to the identification of MSRE sites for the three methylation sequence contexts: *Bsp119I* (TTCGAA) for mCGs, *MspI* (CCGG) for mCHGs, and *HpyF3I* (CTNAG) for mCHHs (**Fig. 6A and B**). An unmethylated D-tDNA^Val^ (AT3G11395) possessing all sites was used as a positive digestion control (**Fig. 6B**).

**Figure 6.**
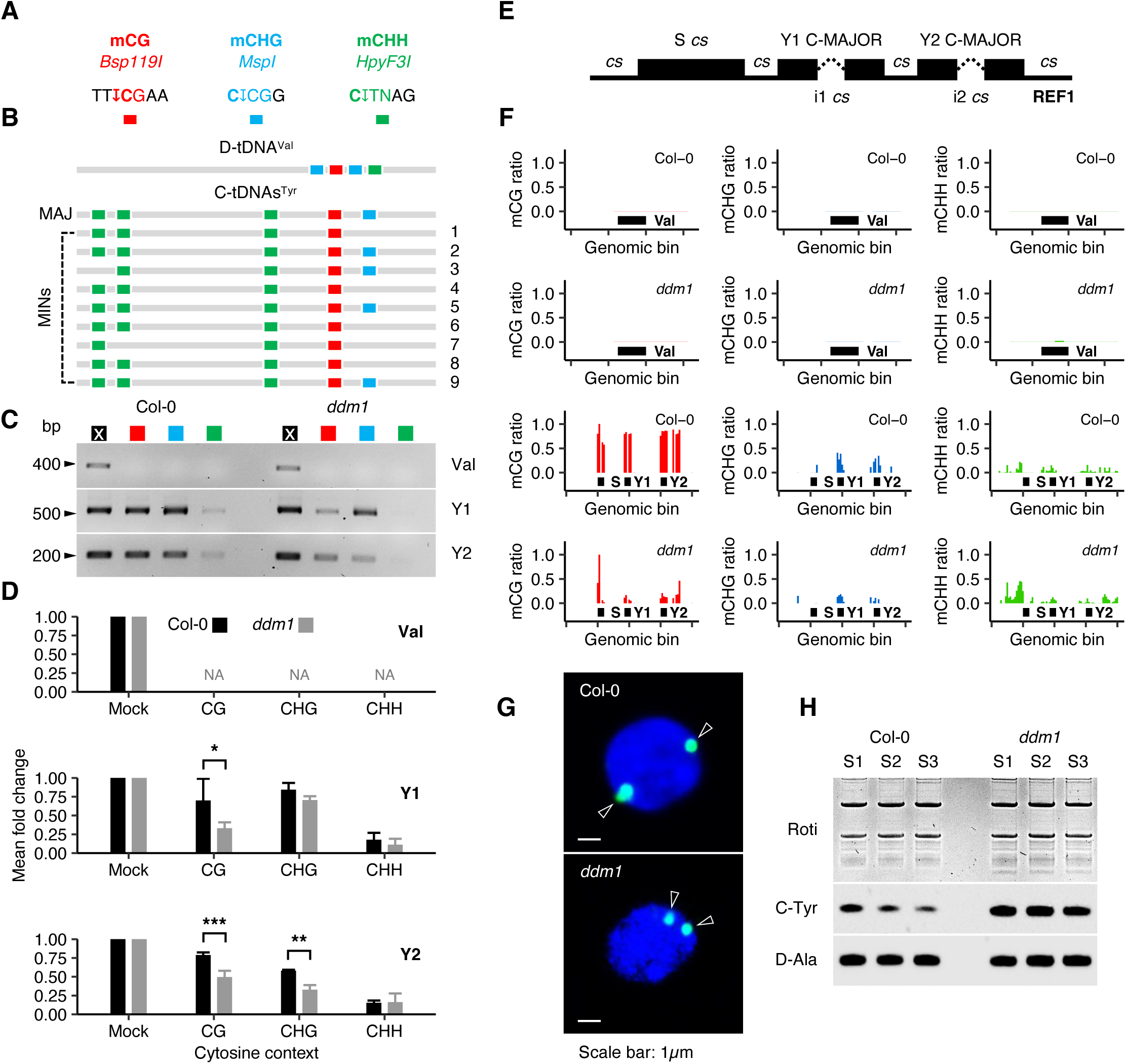
Characterization of SYY cluster chromatin in *ddm1* roots. **A**, Association between the methylation context, chosen restriction enzyme and MSRE site. The cytosine responsible for the methyl-sensitivity is in bold. The cutting position is showed by an arrow. **B**, MSRE site composition of the D-tDNA^Val^ control, major (MAJ) and minor (MIN) C-tDNA^Tyr^ sequences. **C**, Chop-PCR amplification of Val, Y1, and Y2 loci with Col-0 and *ddm1* total root genomic DNA. A cross designates mock treatment. Otherwise, the colour code is similar to the A panel. **D**, Relative chop-PCR product quantities by genotype, gene, and digested cytosine context. Mock is arbitrarily set to 1. Histograms represent the mean of three independent biological replicates, and bars the standard deviations. A two-way ANOVA test disclosed a significant difference in Y1 products accumulation upon *Bsp119I* digestion, and Y2 ones upon *Bsp119I* and *MspI* digestion between Col-0 and *ddm1* (P < 0.05: *, P < 0.01: **, P < 0.001: ***). **E**, Design of NGS data mapping REF1. i: intron. *cs*: consensus. **F**, Methylation levels at S, Y1 and Y2 loci in Col-0 and *ddm1* total root genomic DNA. The D-tDNA^Val^ is presented as control. Bars represent the ratio of methylated cytosines in 20bps bins. **G**, Subnuclear location of the SYY cluster in Col-0 and *ddm1* root nuclei. BAC-FISH signals are identified with arrows. **H**, Northern blot detection of C-tRNAs^Tyr^ and D-tRNAs^Ala^ in the total RNA extracts of Col-0 and *ddm1* root sections. S1: mitosis + tip, S2: transition zone, S3: elongation zone.

The Val control is not amplified in any experimental context (**Fig. 6C and D**). It ascertains that this locus is unmethylated, and that input DNA is fully digested. By contrast, wild-type Y1 and Y2 loci resist *Bsp119I* and *MspI* digestions and were amplified (**Fig. 6C and D**). It confirms that these genes are predominantly CG and CHG methylated in Col-0 roots. However, Col-0 Y1 and Y2 genes did not resist *HpyF3I* digestion and were only weakly amplified (**Fig. 6C and D**). It shows that they are only barely methylated in the CHH context, that is a hallmark of the *A. thaliana* methylation landscape (Law and Jacobsen, 2010; Zhang et al., 2018). Chop-PCR amplifications using *ddm1* template DNA revealed enhanced Y1 and Y2 sensitivities to digestions. It substantiated that the MSRE design detects significant DNA hypomethylations in CG (Y1 and 2) and in CHG (Y2) contexts (**Fig. 6C and D**). Y1 amplifications remain high and persistent for the CHG context whatever the genotype (**Fig. 6C and D**). It is due to 12 MINORs not holding the *MspI* site and mainly standing at the Y1 position (**Figs. 1 and 6C-D**). No significant changes in mCHH levels were observed between Col-0 and *ddm1* with this approach (**Fig. 6C and D**).

To verify that DNA methylation dynamics at chosen chop-PCR sites reflect what happens for the entire cassette, Col-0 and *ddm1* root methylomes were reanalysed (Zemach et al., 2013). Because the SYY cluster is highly repetitive, TAIR10 reference would undeniably bias the interpretation of DNA methylation levels at genes with identical sequences. To bypass the issue, we edited a new reference REF1 wherein the SYY cluster was replaced by a consensus sequence (**Figs. 6E and S3**). In this, each position represented by more than one nucleotide upon alignment was considered as degenerate (N). Y1 and Y2 tDNA sequences were both blocked with the MAJOR to correlate DNA methylation dynamics with northern blot analysis. Introns i1 and i2 were however kept consensual and are by chance different (**Figs. 6E and S3**). Overall, such design allowed us to confidently assess MAJOR methylation status at Y1 and Y2 cassettes.

Our analysis confirmed that the Val control is unmethylated, while S, Y1, and Y2 cassettes are predominantly methylated (mCG > mCHG > mCHH) (**Fig. 6F**). In *ddm1*, CG and CHG hypomethylations were observed, while mCHHs were unchanged (**Fig. 6F**). The hypo-methylation of the SYY cluster does not correlate with a subnuclear relocation of this genomic region in root 2C nuclei (**Fig. 6G**). Indeed, irrespective of Col-0 or *ddm1*, FISH signals were systematically observed close to the nuclear periphery (**Fig. 6G**). Nevertheless, the expression pattern of the hypomethylated SYY cluster is altered among the three main root developmental zones *i.e.,* the tip and division zone (S1), transition zone (S2), and elongation zone (S3) (**Fig. 6H**). In accordance with **Fig. 2C-D**, the amount of C-tRNAs^Tyr^ gradually decreases in Col-0 from S1 to S3 total RNA extracts (**Fig. 6H**). However, their detection becomes strong and homogenous in corresponding *ddm1* extracts (**Fig. 6H**).

Surprisingly, the two methods established that the gradient expression of the SYY cluster in Col-0 roots occurs despite of no significant changes in DNA methylation levels among the three segments S1-3 (**Fig. 7A and B**). The results contrast with the strong and homogenous expression of the transgenic 1xSYY repeat in the absence of DNA methylation (**Figs. 5F and 7C**). The reanalysis of published methylomes also showed no DNA methylation losses in the columella compared to whole tip data (**Fig. 7D**) (Kawakatsu et al., 2016). Instead, high C-tRNAs^Tyr^ accumulation in this tissue correlates with a *de novo* methylation of S, Y1 and Y2 cassettes caused by the RNA-directed DNA methylation pathway (RdDM) (**Fig. 7D**) (Kawakatsu et al., 2016; Zhang et al., 2018).

**Figure 7.**
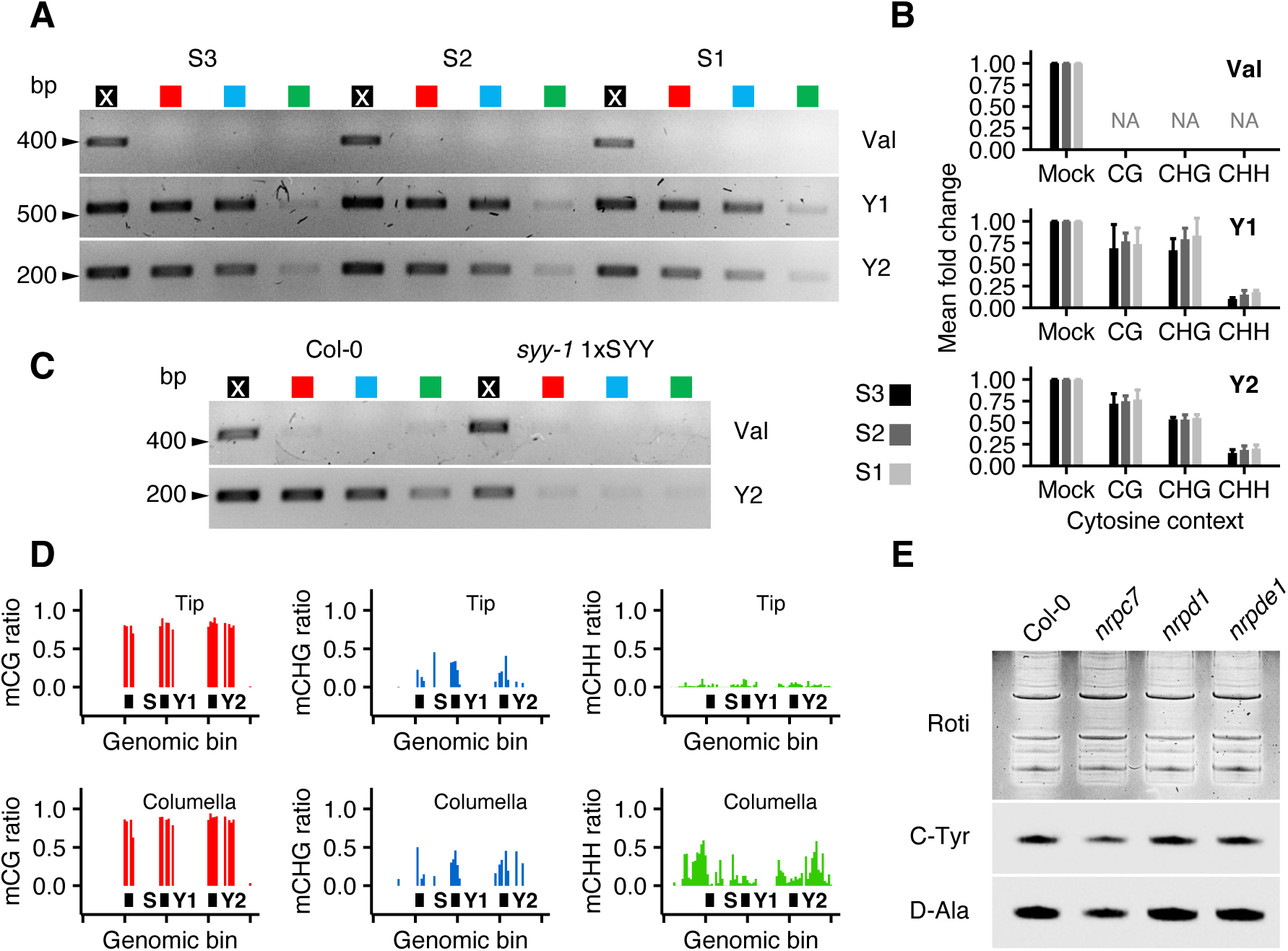
Methylation landscape of natural and transgenic SYY repeats in roots. **A**, Chop-PCR amplification of Val, Y1 and Y2 loci with Col-0 root S1, S2 and S3 genomic DNA. **B**, Relative chop-PCR product amounts by gene, digested cytosine context and root segment. Mock quantification value is arbitrarily set to 1. Histograms represent the mean of three independent biological replicates, and bars the standard deviation. **C**, Chop-PCR amplification of Val and Y2 loci with Col-0 and *syy-1* 1xSYY total root genomic DNA. **D**, Methylation levels at S, Y1 and Y2 loci in whole tip and columella genomic DNA. Bars represent the ratio of methylated cytosines in 20bps bins. **E**, Northern blot detection of C-tRNAs^Tyr^ and D-tRNAs^Ala^ in the total RNA extracts of Col-0, *nrpc7*, *nrpd1* and *nrpde1* root S1.

Altogether, these results demonstrate that DNA methylation has a role in the developmental regulation of the SYY cluster along root. The analysis of the *ddm1* mutant further established the negative impact of this epigenetic mark on SYY cluster expression, and revealed that its transcription is possible at the nuclear periphery when hypomethylated. Surprisingly, the natural expression of the SYY cluster is established along roots via a classical RNAPIII transcriptional system, and despite elevated DNA methylation levels. Indeed, a partial loss-of-function mutant for a RNAPIII-specific subunit (*i.e., nrpc7*) leads to decreased C-tRNAs^Tyr^ and D-tRNAs^Ala^ in S1 total RNA extracts (**Fig. 7E**). By contrast, knock-out mutants for RNAPIV– and V-specific subunits (*i.e., nrpd1* and *nrpde1*) were used as negative controls, and have no effects on C-tRNAs^Tyr^ and control D-tRNAs^Ala^ accumulations in S1 total RNA extracts (**Fig. 7E**).

### Root tip cells are compatible with Ser, Tyr, and Pro-rich translation

Correlating amino acid frequencies in gene coding sequences with tDNA copy numbers showed that C-tDNAs are excessive copies (Michaud et al., 2011; Hummel et al., 2020). It signifies that the expression of D-tDNAs theoretically produces tRNA amounts sufficient to translate the putative mRNA pool encoded in the *A. thaliana* genome.

Therefore, the existence of excessive tDNAs, for instance those in the SYY cluster, might hint a translatome demanding on high levels of the corresponding Ser and Tyr in particular cells or developmental stages. Following this logic, we datamined *A. thaliana* genome for proteins rich in these amino acids. The analysis pointed out a subset of proteins outlying *A. thaliana* proteome and co-enriched in Ser and Tyr (**Fig. 8A**). EXTENSINs (EXTs) 22, 6, 9, 18 and 19 are the top five (**Fig. 8A and B**). EXTs belong to a larger superfamily (*c.a.* 166 genes) of cell wall and secreted components named hydroxyproline-rich glycoproteins or HRGPs (Showalter, 1993; Showalter et al., 2010; Hijazi et al., 2014; Castilleux et al., 2018; Driouich et al., 2021; Moussu and Ingram, 2023). The primary sequence of dicot EXTs entails tandem repetition of a low-complexity motif containing stretches of Pro plus Ser and Tyr residues (**Fig. 8B**) (Showalter, 1993; Showalter et al., 2010). The three amino acids can represent up to 81% of the total protein content. Interestingly, the co-enrichment of Pro with Ser and Tyr also correlates with the existence of tDNA^Pro^ clusters in *A. thaliana* nuclear genome (Hummel et al., 2020; Hummel and Liu, 2022).

**Figure 8.**
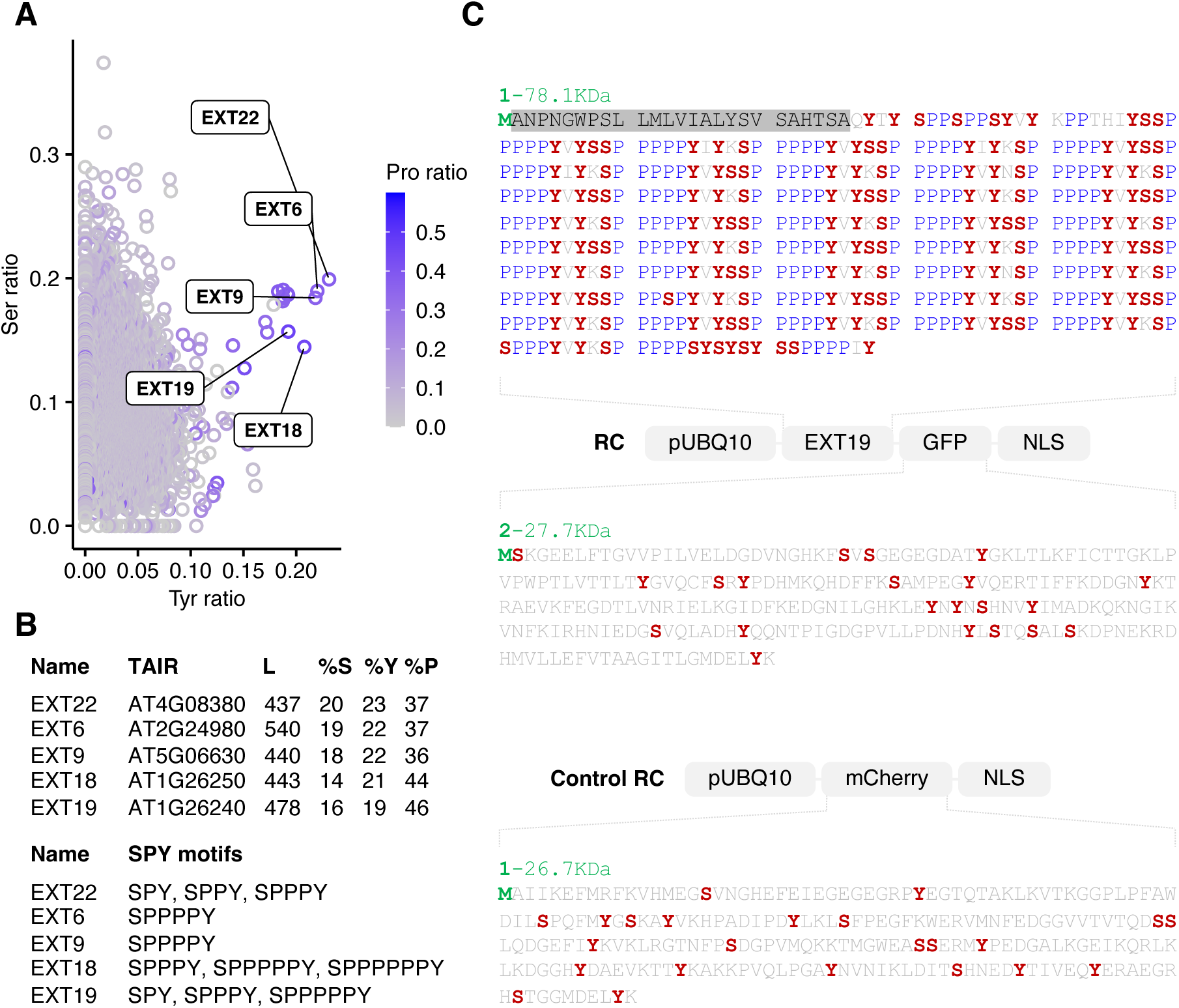
Design of a reporter system to monitor Ser, Tyr and Pro-rich translation. **A**, Identification of proteins co-enriched in Ser and Tyr encoded in the *A. thaliana* Col-0 genome. A colour gradient informs about the Pro content of individual proteins. Names of the top five proteins are flagged. **B**, Listing of EXT names, TAIR IDs, lengths (L), amino acid percentages (%) and SP_n_Y motifs. **C**, Reporter constructs (RCs) used to study Ser, Tyr and Pro-rich translation *in planta*. In protein sequences, initiator Met residues are in green, Ser and Tyr ones in red, and Pro stretches in blue. The putative molecular weight of translational products is indicated in green. The removed EXT19 secretion motif is highlighted in grey.

To monitor high Ser, Tyr and Pro translational demands *in planta*, we designed a reporter construct (RC) forcing the strong and constitutive expression of the EXT19 coding sequence fused to the GFP (**Fig. 8C**). The constitutive UBIQUITIN10 promoter and its intron-containing 5’UTR were used aiming at a constant transgene expression in all tissues. (Norris et al., 1993; Sun and Callis, 1997) (**Fig. 8C**). To hinder the integration of the heterologous protein in the cell wall, the putative N-ter secretion signal of the EXT19 was removed and antagonized by a NLS tag in C-ter of the construct (**Fig. 8C**). The cytoplasmic retention of the protein was preferred to purify it without highly destructive methods. Indeed, EXTs can be covalently crosslinked with others HRGPs or others cell wall constituents, what makes them insoluble and difficult to extract (Moussu and Ingram, 2023). Also, the cytoplasmic retention was preferred in case EXT19 is primarily secreted in the environment. The fluorescent tag was intentionally put in C-ter of the construct, so that the pre-requisite for its accumulation is a prior translation of the Ser, Tyr, and Pro-rich cassette (**Fig. 8C**). We also designed a control RC wherein the EXT19-GFP was replaced by a fluorescent protein not particularly enriched in Ser, Tyr, and Pro *i.e.,* mCherry (**Fig. 8C**).

The RC was stably agrotransformed in the Col-0 background, and the accumulation of the fusion protein was followed at the tissular level in defined germination timepoints (**Fig. 2A**). A strong accumulation of EXT19-GFP-NLS was observed in whole 0DAS embryos (**Fig. 9A**). The developmental stage validated that RC mRNAs can be ubiquitously expressed and translated. Starting from 1DAS and maintained till the adult 14DAS stage, the constitutive accumulation of EXT19-GFP-NLS in roots is clear in favour of a cell-specific expression in the four peripheral RCC and adjacent LRC cells (**Fig. 9A**). Interestingly, protein level progressively increases from the first layer of cells (immediately below the meristem) to the peripheral one in detachment (**Fig. 9A**). Thus, RC expression follows a spatiotemporal logic in these tissues related to the maturation and specialization of RCC and LRC cells. Parallelly, EXT19-GFP-NLS accumulation gradually decreases with cotyledons growth, and was not detected in adult leaves and in untransformed WT plants (**Fig. 9A**). The co-agrotransformation of the RC and control RC ascertained that the EXT19-GFP protein is not nuclear despite the presence of a NLS tag, but rather accumulate in dense cytoplasmic foci which can aggregate around nuclei (**Fig. 9B**). It might result from a conflict between others secretion signals in the protein sequence and its nuclear import.

**Figure 9.**
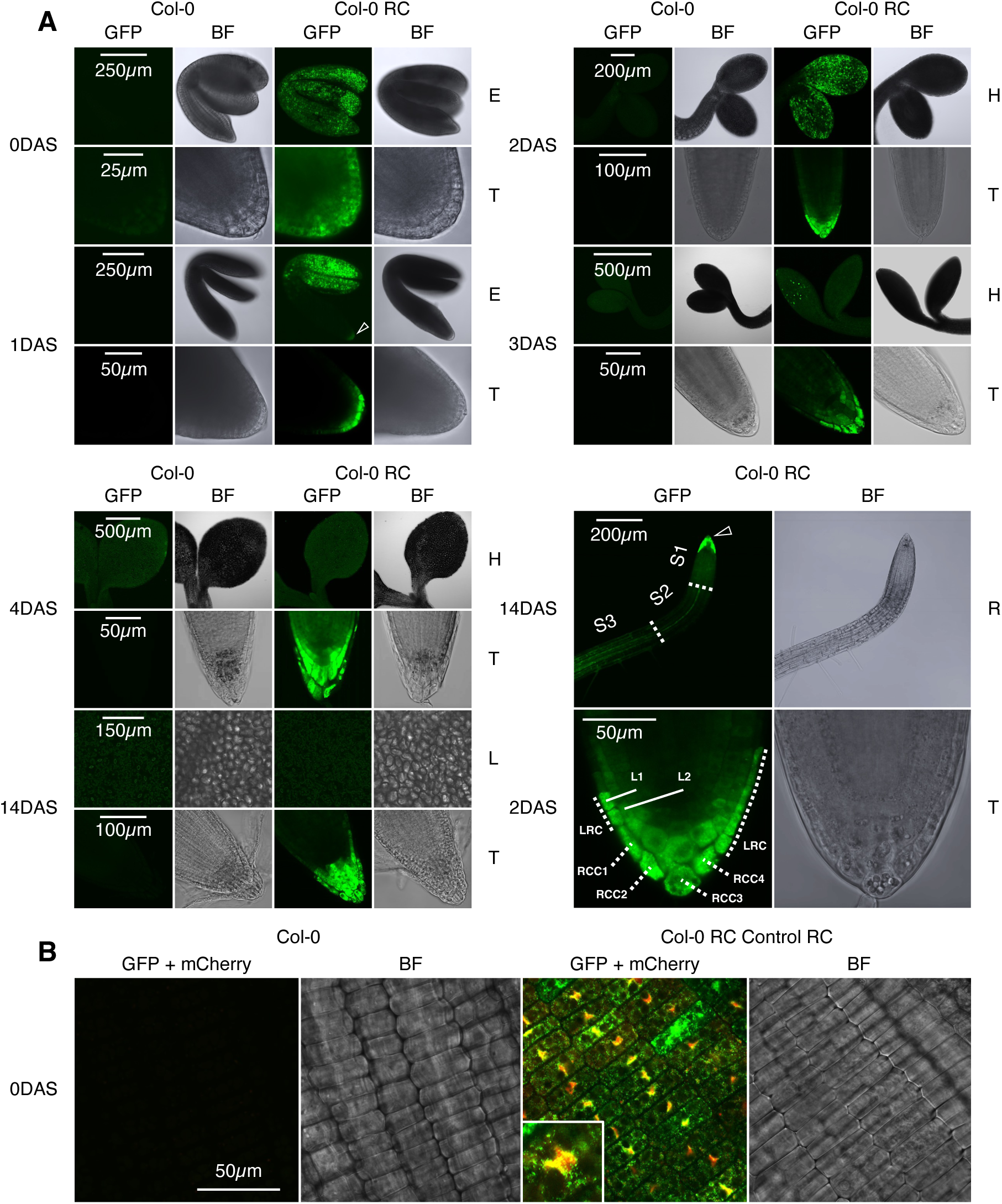
Tissular and subcellular accumulation of reporter proteins. **A**, Representative confocal snapshots of 0, 1, 2, 3, 4, and 14DAS Col-0 and Col-0 RC plants. The white arrow points out the apparition of the RCC and LRC-specific fluorescence. E: embryo, T: tip, R: root, H: hypocotyl, L: leaf. **B**, Representative confocal captions of 0DAS Col-0 and Col-0 RC Control RC radicle cells. A magnification on a single nucleus is shown.

Then, we performed a genetic cross to bring the RC and control RC into *syy-1* plants and examine how EXT19-GFP-NLS and mCherry-NLS synthesis can be influenced by the absence of the SYY cluster. Concomitant with a high proportion of Ser and Tyr codons in the RC, we found a significant decrease of GFP fluorescence in RCC and LRC upon expression in the *syy-1* background compared to Col-0 plants (**Fig. 10A and B**). The coexpression of the RC and control RC in *syy-1* led to a similar effect, but mCherry fluorescence remained unaffected in RCC/LRC and surrounding tissues compared to Col-0 (**Fig. 10C-E**). An indicator of fluorescence intensity revealed that, under the dependence of pUBQ10, the mCherry-NLS is overexpressed in an internal cluster of cells (**Fig. 10D**). This profile does not overlap with the GFP pattern, further confirming that the accumulation of the Ser, Tyr, and Pro-rich reporter in RCC and LRC cells is not the consequence of the pUBQ10 activity.

**Figure 10.**
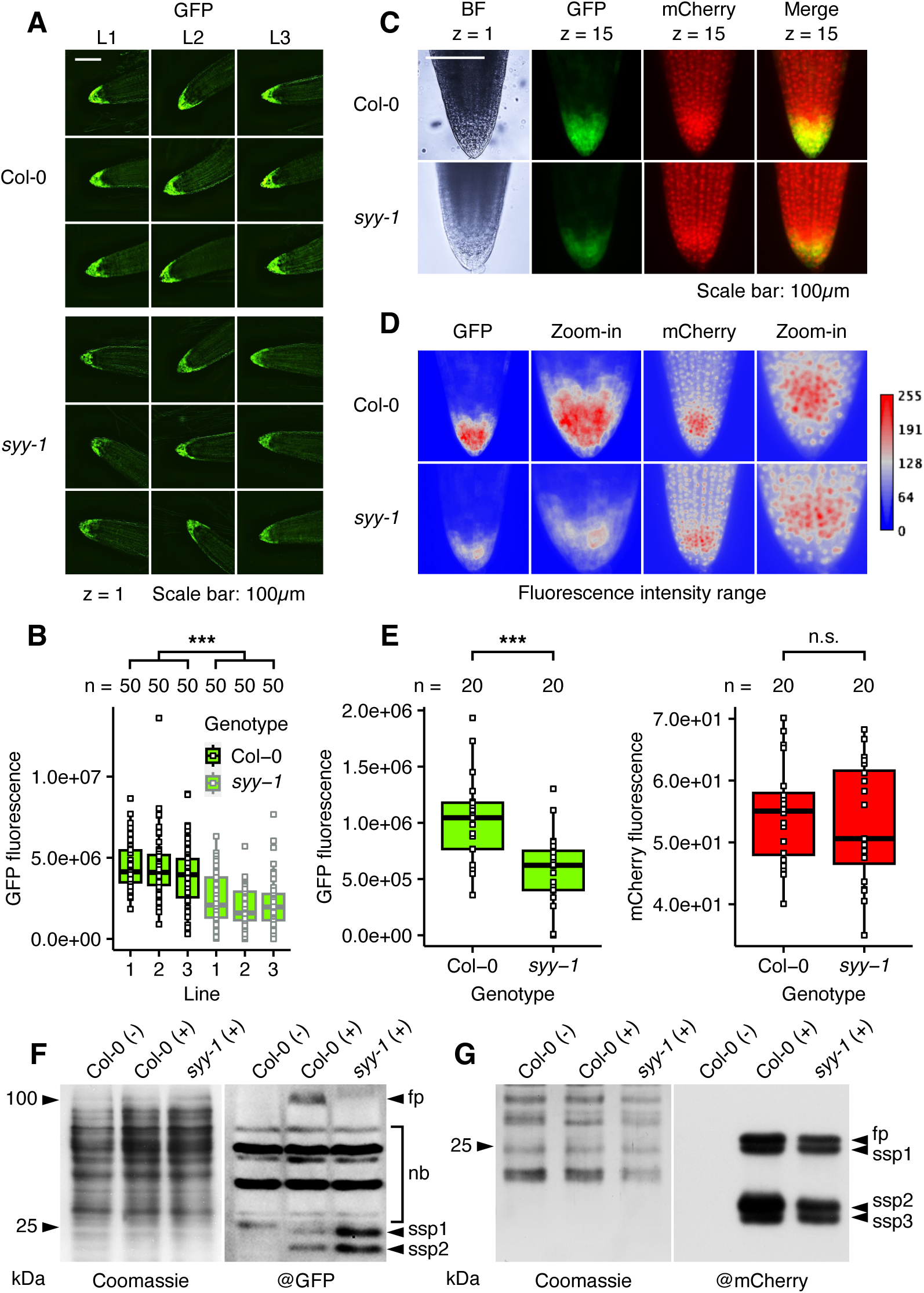
Accumulation of reporter proteins in Col-0 and *syy-1* backgrounds. **A**, Analysis of EXT19-GFP-NLS accumulation in Col-0 and *syy-1* tips by epifluorescence (one focal plan z). Three independent lines (L) are shown by genotype. For each, three representatives are depicted. **B**, Distribution of GFP fluorescence intensity by genotype and line. 50 individuals (n) were analysed by line. A one-way ANOVA test disclosed a significant difference in GFP fluorescence between Col-0 and *syy-1* populations (P < 2×10^−16^, ***). Thumbnails in A were selected based on a homogenous dispersal around medians plotted in B (*i.e.,* +/− 500000). **C**, Analysis of EXT19-GFP-NLS and mCherry-NLS co-accumulation in Col-0 and *syy-1* tips by epifluorescence (merge of 15 focal plans). BF: bright field. **D**, Range indicator of GFP and mCherry fluorescence intensity in Col-0 and *syy-1* tips. **E**, Distribution of GFP and mCherry fluorescence intensity by genotype. 20 individuals (n) were analysed by line. A one-way ANOVA test disclosed a significant difference in GFP fluorescence between Col-0 and *syy-1* populations (P = 0.00034, ***) while no significant difference (n.s.) was observed for mCherry fluorescence. Thumbnails in C were selected based on a homogenous dispersal around medians plotted in E (*i.e.,* +/− 150000). **F**, Immunodetection of EXT19-GFP-NLS in the total root extracts of Col-0 and *syy-1* individuals expressing (+) or not (-) the RC. **G**, Immunodetection of mCherry-NLS in the total plant extracts of Col-0 and *syy-1* individuals expressing (+) or not (-) the control RC. fp: full protein, nb: nonspecific bands, ssp: subspecies.

EXT19-GFP-NLS and mCherry-NLS accumulations were monitored in Col-0 and *syy-1* RC plants using western blots (**Fig. 10F and G**). A signal around 100kDa, corresponding to EXT19-GFP-NLS, was specifically detected in RC lines (**Fig. 10F**). Downregulated in *syy-1* protein extract, its behaviour reflects what observed for the root tip fluorescence (**Fig. 10A-E**). Surprisingly, an enhanced accumulation of shorter protein subspecies is detected in the *syy-1* extract (**Fig. 10F**). They might correspond to degradation products reflecting an impaired proteostasis. Importantly, the presence of these subspecies further confirms that the difference in the EXT19-GFP-NLS accumulation between Col-0 and *syy-1* is not due to an RNA silencing mechanism. The blot also suggests that *syy-1* root tip fluorescence might only be explained by partial GFP fragments with altered emission efficiency. By contrast, SYY cluster deletion has no impact on the accumulation of the control mCherry-NLS and its subspecies (**Fig. 10G**).

Altogether, our data show that the homeostasis of RCC and LRC tissues is compatible with a continuous translation and stabilization of an overexpressed EXTENSIN along seedling establishment. It hints that these tissues are physiologically equipped to elaborate Ser, Tyr, and Pro-rich translatomes. RCC and LRC tissues host the high expression of the SYY cluster, and its deletion alters the accumulation of the EXTENSIN reporter. It evidences that this genomic region is involved in this tissular peculiarity via the production of tRNA amounts favouring the context-dependant translation of SP_n_Y motifs.

## DISCUSSION

Hereinabove, we support that a subset of tRNA^Tyr^ genes, tandemly organized with tRNA^Ser^ ones in a genomic structure called SYY cluster, is subject to a developmental expression pattern along the root (**Figs. 1-6**). Cell layers bordering protective tissues of the meristem *i.e.,* the columella and lateral root cap (Dolan et al., 1993) are the main transcriptional hotspots of the SYY cluster during seedling establishment (**Fig. 4**). This study strengthens growing experimental evidence that the tRNAome is highly dynamic. Only few of the redundant tRNA genes inserted in a nuclear genome are expressed in a given condition, and the unequal contribution is dynamically reshaped following developmental and environmental stimuli (Canella et al., 2010; Moqtaderi et al., 2010; Oler et al., 2010; Gogakos et al., 2017; Thornlow et al., 2018; Hummel et al., 2019; Torres, 2019; Torres et al., 2019; Hummel et al., 2020; Liu and Sun, 2021; Ma et al., 2021; Hummel and Liu, 2022; Hughes et al., 2023; Kapur et al., 2024).

In this work, we provide evidence that the gradient expression of the SYY cluster along root occurs despite elevated CG methylation levels (**Figs. 6-7**). Its behaviour strikingly tallies the one of 5S rRNA gene clusters, tandem repeats of which are also transcribed by RNAPIII despite elevated CG methylation levels in *A. thaliana* (Mathieu et al., 2002). These results contrast with the previous work, thereby it was showed that CG methylation is involved in the silencing of clustered tRNA^Tyr^ genes (Hummel et al., 2020). The strong and constitutive expression of the transgenic SYY repeat further confirmed the negative impact of high DNA methylation levels on tRNA gene transcription (**Figs. 5 and 7**). Same conclusions were drawn in *Xenopus laevis* oocytes, where the injection of an artificially methylated tRNA^Lys^ gene inhibited at 80% its transcription (Besser et al., 1990). Transcribing artificially methylated 5S rRNA genes was however doable *in vivo* and *in vitro* (Besser et al., 1990; Mathieu et al., 2002).

tRNA and 5S rRNA gene transcription do not involve the same RNAPIII promoters and pioneering factors. tRNA genes are characterized by a type 2 promoter and relies on sequential interactions between TFIIIC, TFIIIB and RNAPIII (Roeder, 2019). 5S rRNA genes have a type 1 promoter, and require the additional transcription factor TFIIIA sequentially recruiting TFIIIC, TFIIIB and RNAPIII (Roeder, 2019). Therefore, 5S rRNA gene transcriptional insensitivity to DNA methylation might be explained by a more tolerant pre-initiation complex assembly mediated by TFIIIA binding pattern. This hypothesis is however refuted as TFIIIC and B can bind methylated type 2-like promoters, and several tRNA/Alu-related short interspersed nuclear elements (SINEs) are expressed despite high DNA methylation levels in human cells (Varshney et al., 2015). The alteration of their methylation landscape with DNA Methyltransferase 1 (Dnmt1) knockout, or 5-azacytidine treatments, do not induce their expression but reactivates silent protein-coding genes (Varshney et al., 2015). Instead, authors showed that histone methylations (*i.e.,* H3K9me3) primarily inhibit RNAPIII transcription at SINEs (Varshney et al., 2015). In parallel, Sizer and colleagues exemplify several TFIIIC, TFIIIB and RNAPIII-bound tRNA genes located in CpG islands (Sizer et al., 2022). Still, RNAPIII is recruited at methylated tRNA gene repeats during zygotic genome activation in *Xenopus* and mouse (Parasyraki et al., 2024).

Hence, DNA methylation might not be the main epigenetic mark involved in the silencing of the SYY cluster. In *ddm1* and *met1* rosettes, or in 5-azacytidine/zebularine-treated plants, tRNA^Tyr^ quantities produced by 34 clustered loci remain low relative to the expression of 16 dispersed ones (Hummel et al., 2020). Above conditions have no impact on the silencing of tRNA^Pro^ gene clusters (Hummel et al., 2020). While DNA methylation profile at the SYY cluster does not change along the root, nucleosomes compaction and their composition in histone variants might be more dynamic parameters during development (Rosa et al., 2014; Probst et al., 2020; Zhang et al., 2021; Feng et al., 2022; Probst, 2022; Simon and Probst, 2023). Around 150bps, tRNA gene size is subnucleosomal. Therefore, their expression might be particularly influenced by nucleosome positioning and chromatin compaction. Transcribed tRNA genes are generally nucleosome-free (Shukla and Bhargava, 2018). In yeast, antisense RNAPII activity at tRNA genes triggers histone acetylation, and sets a nucleosome-free environment potentializing RNAPIII transcription (Yague-Sanz et al., 2023).

The high clustered tRNAs^Tyr^ accumulation in columella and lateral root cap independently confirms that chromatin characteristics of these tissues enable the expression of repeats (**Fig. 4**) (Kawakatsu et al., 2016). A heterologous DDM1:GFP protein expressed under the control of native promoter accumulates in all root tissues except columella and lateral root cap in *ddm1* background (Kawakatsu et al., 2016). DDM1 acts as a chromatin remodeler displacing linker H1, allowing access to DNA for methyltransferases MET1, CHROMOMETHYLATES 2 and 3, and DOMAINS REARRANGED METHYLTRANSFERASE 2 (Zemach et al., 2013). DDM1 also facilitates the deposition of histone variant H2A.W at heterochromatin (Osakabe et al., 2021; Osakabe et al., 2023). H1 and H2A.W together assist the compaction of constitutive heterochromatin (Bourguet et al., 2021). The double mutant *h1 h2a.w* leads to a strong decompaction, and occasions the *de novo* methylation of heterochromatin via the RdDM pathway (Bourguet et al., 2021). Remarkably, columella cells are characterized by downregulated H1.1, H1.2, H2A.W6 and H2A.W7 mRNA levels, and a genome-wide RdDM-mediated hypermethylation in the CHH context (Kawakatsu et al., 2016). The situation strikingly phenocopies *h1 h2a.w*, and might reflect a globally decondensed nucleosome environment, which is favourable to the SYY cluster transcription despite a preserved CG methylation maintenance.

Our data show that columella and lateral root cap cell physiology authorizes a continuous translation and accumulation of a transgenic Ser, Tyr and Pro-rich EXTENSIN (**Figs. 8-10**). We evidence a perfect match between the expression of the SYY cluster and the protein (**Figs. 4 and 9**). SYY cluster deletion hinders the capability of such cells to produce the EXTENSIN (**Fig. 10**). Polyproline motifs found in the EXT19 sequence are known to stall ribosomes and require a specialized elongation factor (*i.e.,* EF-P/ eIF-5A), thus making their translation highly sensitive to tRNA availability (Peil et al., 2013; Ude et al., 2013; Starosta et al., 2014; Lassak et al., 2016). Thus, tRNA shortage in *syy-1* might occasion impaired translational rates at S and Y residues flanking polyproline motifs in EXT19 and ribosome collisions. These events might trigger transcripts and/or protein degradation (Vind et al., 2020). Our study mirrors the appealing case of *Bombyx mori* silk glands, where specific tRNA^Ala^, tRNA^Gly^ and tRNA^Ser^ species are upregulated in the organ and confer a translational specialization for the high production of fibroin during the secretion phase (Garel et al., 1973; Majima et al., 1975; Siddiqui and Chen, 1975; Chevallier and Garel, 1982).

Yet, *syy-1* and *syy-2* deletion alleles display no macroscopic phenotypes. They germinate, grow, bolt, flower and pod like Col-0 (**Fig. 11A-D**). Moreover, SYY cluster transcription does not seem to have a developmental meaning in RCC and LRC tissues. No obvious alteration in the cellular organization neither desquamation of these tissues was noted (**Fig. 11E-F**). Upon release, RCC and LRC cells acquire a specialized function in producing a mucilaginous structure known as root extracellular trap (RET), and controlling the interactions with soil-dwelling organisms (Driouich et al., 2011; Hawes et al., 2016; Driouich et al., 2019; Castilleux et al., 2021; Hromadová et al., 2021; Fortier et al., 2023). Whether there is any function of the SYY cluster in promoting Ser, Tyr, and Pro-rich translation during this process remains to be fully elucidated.

**Figure 11.**
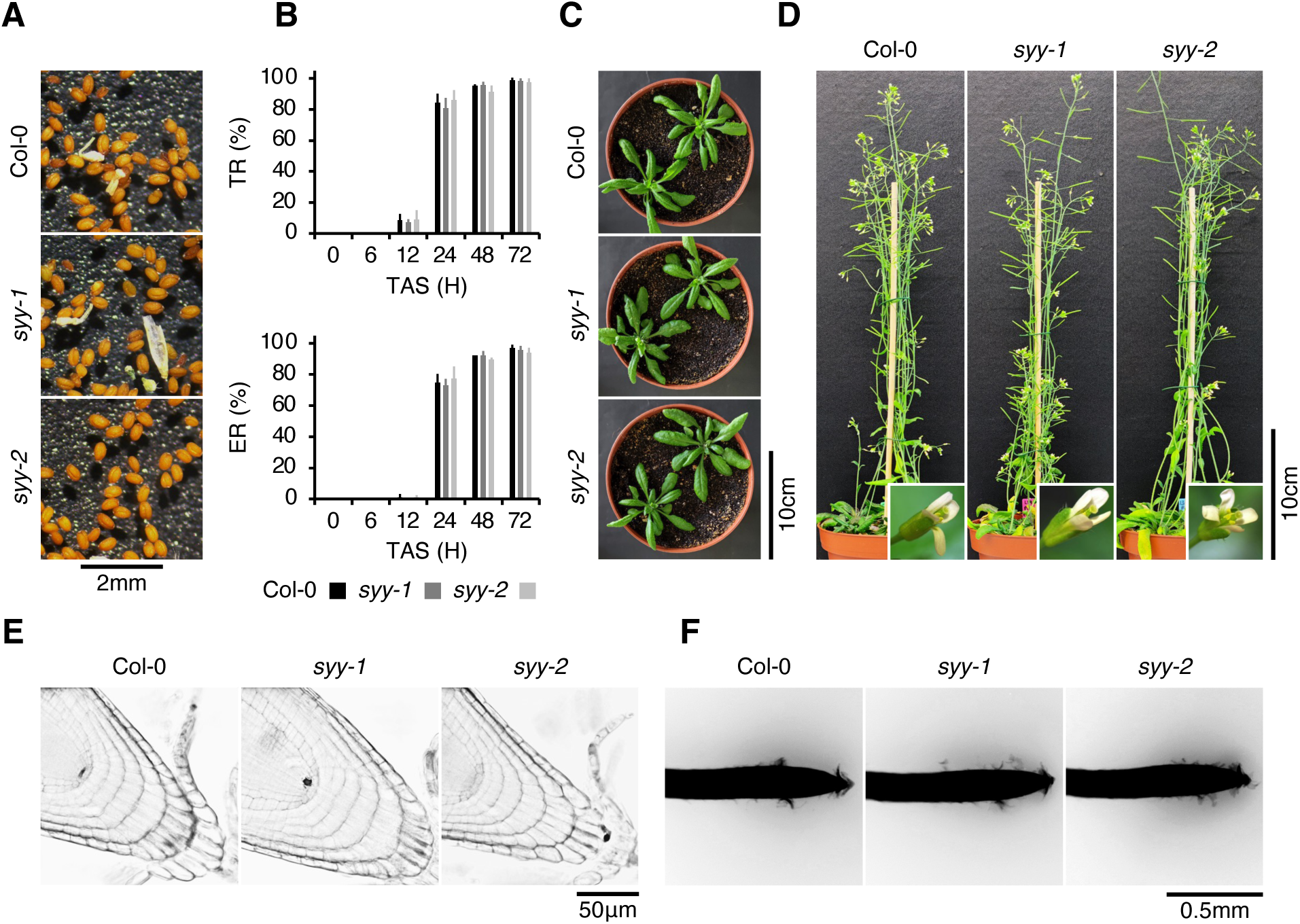
Phenotyping of tRNA gene mutants during development. **A**, Col-0, *syy-1* and *syy-2* dry seeds. **B**, Testa and endosperm rupture kinetics (TR and ER, respectively) of Col-0, *syy-1* and *syy-2* germinating seeds. Histograms correspond to the mean of three independent biological replicates and bars standard deviations. A twoway ANOVA test revealed no significant difference between testa and endosperm rupture kinetics of Col-0, *syy-1* and *syy-2* seeds (P = 0.603 and 0.607, respectively). TAS: time after stratification. **C and D**, Col-0, *syy-1* and *syy-2* rosettes, flowers, and bolted plants with pods. **E**, Propidium ioide staining of 7DAS Col-0, *syy-1* and *syy-2* root tips. **F**, Nigrosin staining of 7DAS Col-0, *syy-1* and *syy-2* root tips.

## METHODS

### Plant material and growth conditions

The following germplasms were used in this study: Col-0 (wild-type), *ddm1-1* (Vongs et al., 1993), *ddm1-2* (Jeddeloh et al., 1999), *ddm1-10* (SALK_093009), *nrpc7-1* (Johnson et al., 2016), *nrpd1-3* (SALK _128428), and *nrpe1-11* (SALK_029919). Seeds were surface-sterilized for 15min in 70% ethanol, 1% Tween-20, and twice for 2min in 100% ethanol. They were sown on a half-strength Murashige and Skoog medium pH5.7 (M0255, Duchefa Biochemie) supplemented with 1% sucrose and 0.6% phytagel (Sigma-Aldrich). Seeds were stratified for 48h at 4°C, and vertically grown at 21°C on a CS 250/300_5_3 (Photon Systems Instruments) cultivation shelf equipped with LEDs (intensity: 100µmol.m^−2^.s^−1^, photoperiod: 16h).

### Total genomic DNA extraction and sequencing

gDNA was prepared according to (Michiels et al., 2003). Sequencing of vectors, and deletion zone PCR products was performed by Eurofins Genomics.

### Cloning

Dual and multiplexed CRISPR/Cas9 plasmids were obtained as described in (Zhang et al., 2016; Wu et al., 2018). To obtain the Ser, Tyr and Pro-rich reporter construct, the UBIQUITIN10 (AT4G05320) promoter and intron-containing 5’UTR, and the EXTENSIN19 (AT1G26240) coding sequence excluding the N-terminal secretion motif and stop codon were amplified from Col-0 gDNA. They were Gibson assembled, together with a GFP-NLS cassette, in a pGREEN-IIS destination vector (Hellens et al., 2000) carrying the Rubisco small subunit (rbcs) terminator, and conferring spectinomycin (bacteria) and Basta (plant) resistances (see pFK-206 in MPI for Biology Tübingen vector list). A double glycine spacer was placed between EXT19 and GFP-NLS genes. The control reporter construct was obtained identically, except that the EXT19-GFP-NLS region was replaced by a mCherry-NLS. The engineered tRNA^Tyr^ locus was produced as gene strands by Eurofins Genomics, and the SYY cluster repeat was amplified from Col-0 gDNA. They were individually cloned into a pFK-206 plasmid by Gibson assembly. A mCherry gene expressed under the control of the seed-specific 2S3 promoter was used as selection marker.

### Southern blotting

The methodology described in (Brown, 1993) was essentially applied, with minor adjustments. For each sample, 2.5µg of gDNA were digested overnight and separated on a 1% agarose gel subsequently treated with depurination and neutralization buffers. Then, gDNA was transferred on a positively charged nylon membrane (Amersham Hybond-N+, GE Healthcare) with a vacuum transfer system (BIO-RAD, Model 785, Vacuum Blotter), and crosslinked with a UVP Crosslinker (AnalytikJena) using following parameters:

Energy-1200 (*100µJ/cm2), twice. A mCherry-specific probe was amplified from a plasmid containing the transgene of interest, and digoxigenin-labelled with DIG High Prime DNA Labelling and Detection Starter Kit II (Roche). Hybridization was performed overnight at 55°C in hybridization buffer (DIG Easy Hyb, Bottle 7, Roche), followed by stringent washes twice in 2x SSC, 0.1% SDS for 15min at RT, and twice in 0.5x SSC, 0.1% SDS for 15min at 55°C. Membrane was then incubated in blocking buffer (DIG Easy Hyb, Bottle 6, Roche) for 30min at RT, and further incubated for 30min at RT in blocking buffer containing Anti-DIG AP conjugate (1:1000, DIG Easy Hyb, Roche). Probes were detected with CSPD substrate (DIG Easy Hyb, Bottle 5, Roche), and with iBrightTM CL750 Imaging System.

### Semi-quantitative chop-PCR

For each reaction, 10ng of gDNA were digested for 30min at 37°C in a 10µl mix containing 0.2µl of either Fast Digest (FD) *Bsp119I*, *MspI*, *HpyF3I* (Thermo Scientific), and 1µl of appropriate buffer. Reactions were inactivated for 20min at 80°C, and digested gDNA was isolated with 5V of SPRI beads (Beckman Coulter). After a 10min incubation, beads were sedimented with a magnetic stand, and supernatants removed. Beads were cleaned twice in 80% ethanol and air-dried. Elution was done in 10µl of 10nM Tris pH8.0. For each PCR amplification, 1.5µl were used as template with the ALLin™ HiFi DNA Polymerase kit (HighQu). Then, 2µl of 6x DNA loading dye (Thermo Scientific) were added, and samples were run along with a GeneRuler 1kb plus DNA ladder (Thermo Scientific), in a 1% agarose gel containing 0.05µl/ml ROTI GelStain (Carl Roth). Amplification products were analysed with a Vilber QUANTUM CX5.

### Isolation of nuclei and fluorescent *in situ* hybridization

Total nuclei were extracted and sorted using a S3e Cell Sorter (Bio-Rad) as delineated in (Zhu et al., 2017; Wang et al., 2021). FISH experiments were realized with 2C nuclei, and the BAC F9K23 (GenBank: AC082643) as explained in (Montgomery et al., 2022).

### Total RNA extraction

For germination time-courses, a protocol yielding in high-quality total RNAs (Chang et al., 1993) was used. For others experiments, RNAs were directly TRIzol-chloroform extracted. Concentration and quality were determined with a DeNovix DS-11 FX+.

### RNA sequencing and data processing

Total RNAs were extracted from 7DAS seedlings using the RNeasy Plant Mini Kit (Qiagen). Following DNaseI treatment (Thermo Scientific, Waltham, MA, USA), total RNAs were incubated with first-strand buffer and subjected to thermal cycling at 80°C for 2min, followed by 94°C for 1.5min. Polyadenylated RNAs were then isolated from total RNAs and processed according to manufacturer’s instructions (NEBNext^®^ Ultra™ II RNA Library Prep Kit for Illumina^®^). After end-repair of double-stranded cDNAs, free ends were ligated to reverse adapters, followed by final enrichment via PCR to construct sequencing libraries.

### Northern blotting

RNAs were routinely loaded in a gel consisting of 15% acrylamide/bis-acrylamide (19/1), 7M urea and 1x TBE, and resolved by electrophoresis at 150V constant in 1x TBE. RNA integrity was checked with ROTI GelStain. Then, RNAs were electrotransfered onto a Hybond-N+ nylon membrane (Amersham) for 75min at 300mA constant and 4°C in 0.5x TBE. RNA crosslinking was performed with an UVLink 1000 Crosslinker (AnalytikJena, energy mode, 2 x 120mJ/cm^2^). Probes were hybridized overnight at 55°C in 6x SSC, 0.5% SDS, and at a final concentration of 20nM. Washings were performed twice in 2x SSC for 10min at 55°C, once in 2x SSC, 0.1% SDS for 30min at 55°C, and for 10min at 25°C. Then, the membrane was rotated for 1h at 25°C in 2x SSC, 0.1% SDS supplemented with 1:15000 (v/v) HRP-Streptavidin (Abcam), and washed three times in 2x SSC, 0.1% SDS for 10min at 25°C. Probes were detected with sensitive HRP substrates (Takara), and an iBrightTM CL750 Imaging System.

### Whole-mount *in situ* hybridization coupled to tyramide signal amplification

The WISH procedure was essentially executed as described in (Wójcik et al., 2018), with adjustments. The material was fixed for 3h in 3.7% formaldehyde, and permeabilized for 90min at 37°C in 75µg/ml proteinase K. The material was hybridized for 16h at 37°C with 100nM biotinylated probe, and washed three times for 10min at 37°C in 2x SSC, then twice for 30min at 42°C in 2x SSC, 0.1% SDS, and three times for 10min in 1x PBS. For the TSA procedure, endogenous peroxidases were first quenched by incubating the material for 1h in a peroxidase suppressor solution (Thermo Scientific). After six baths of 5min in 1x PBS, and one of 5min in 1x PBS, 0.1% Tween 20 (PBST) at 25°C, samples were incubated for 1h at 25°C in 1x PBST, 1:15000 (v/v) HRP-Streptavidin. Upon six baths of 5min in 1x PBST, samples were incubated for 1h in 1:50 of a digoxigenin-tyramide signal amplification solution (TSA Plus DIG, PerkinElmer) prepared according to manufacturer instructions. After being washed six times for 5min in 1x PBST, and once for 30min in 1x PBST, 5% BSA, samples were incubated overnight at 4°C in 1x PBST, 5% BSA, 1:2500 anti-digoxigenin antibodies fused to the alkaline phosphatase (Roche). Washings were performed twice for 30min in 1x PBST, 5% BSA and three times for 10min in 1x PBST. Samples were then incubated three times for 5min with 1x TNM buffer [100mM Tris pH9.5, 100mM NaCl, 50mM MgCl_2_], and staining was performed in 1x TNM supplied with 1:50 NBT/BCIP (Roche). The reaction was finally quenched with three baths of 5min in 1x TE buffer. For the detection of C-tRNAs^Tyr^ in complemented *syy-1* Ing-Y and *syy-1* 1xSYY lines, a regular WISH procedure was executed as described above with a digoxigenin-labelled probe.

### Western blotting

Proteins were TRIzol-chloroform extracted and resuspended in a buffer containing 4.4M urea, 11% glycerol, 88mM Tris-HCl pH6.8, 2.2% SDS, and supplied with 10mM DTT. Proteins were stacked in 6% acrylamide/bis-acrylamide (37.5/1), 125mM Tris-HCl pH6.8 and 0.1% SDS, then resolved in 12% acrylamide/bis-acrylamide, 380mM Tris-HCl pH8.8 and 0.1% SDS. Electrophoresis was performed at 30mA constant, in 25mM Tris, 200mM glycine and 0.1% SDS. Proteins were electrotransfered onto a PVDF membrane (Immobilon-P) preactivated in methanol, at 4°C and 80mA constant for 45min. Proteins were stained in 0.1% Coomassie blue, 7% acetic acid, 50% methanol, washed in 7% acetic acid, 50% methanol and pure methanol. The membrane was blocked overnight at 4°C in TBS-Tween 0,2% (TBST) supplied with 5% milk. Primary antibody incubation was realized for 1h with 1:5000 (v/v) anti-GFP antibody (ab290, Abcam). The membrane was washed in TBST 5% milk, three times for 10min. Secondary antibody incubation was performed for 30min with 1:10000 (v/v) anti-rabbit IgG-peroxidase antibody (Sigma-Aldrich). Again, the membrane was washed three times for 10min in TBST 5% milk, then three times for 10min in TBST. HRP substrates (Takara) and an iBrightTM CL750 Imaging System were used for signal acquisition. For mCherry immunodetection, proteins were resolved in a 10% SDS-PAGE, and electrotransfered onto an Immobilon-P membrane (Millipore). Primary and secondary incubations were performed with a 1:5000 (v/v) 6G6 anti-RFP antibody (ChromoTek), and a 1:10000 (v/v) anti-mouse IgG-peroxidase antibody (Invitrogen) respectively. Super RX medical X-ray films (Fujifilm) were used for signal acquisition.

### Bioinformatics

tRNA sequences and intron informations were retrieved from the PlantRNA 2.0 database (Cognat et al., 2022). tRNA structures were predicted using R2DT (Sweeney et al., 2021). Bisulfite datasets were analysed as described in (Zhu et al., 2017; Hu et al., 2019), except that the reads mapping was conducted with a modified TAIR10 genome reference wherein the SYY cluster was replaced by one consensual SYY repeat sequence (**Fig. S3**). Bisulfite files of roots: SRR578936, SRR578937, SRR578926, SRR578927. Bisulfite files of root tissues: SRR3311822, SRR3311825. RNA-seq reads were aligned against the TAIR10 annotations using TopHat 2 (v2.1.1) with default parameters. The track files of individual RNA-seq samples in bigWig format and the count table were generated with the bamCoverage function from deepTools and the R package GenomicAlignments, respectively (Kim et al., 2013; Lawrence et al., 2013; Ramírez et al., 2016).

### Microscopy and image analysis

Root tip cell organization was prospected in 10μg/mL propidium ioide (PI), and images were acquired with an Olympus IX83 microscope supplied with an Olympus XM10 camera and the CellSens Dimension v1.11 software (Olympus). Tip cell release was analysed in 5% Nigrosin (Sigma-Aldrich), and images acquired with a KERN OPTICS OZL 464 stereo microscope supplied with a tablet Cam ODC 241 (KERN). EXT19-GFP-NLS accumulation pattern was captured using an LSM880 confocal microscope (Zeiss) and ZEN black v2.3 software. FISH results were screenshot with an LSM700 confocal microscope. EXT19-GFP-NLS and mCherry-NLS accumulation in Col-0 and *syy-1* backgrounds was analysed similarly to PI experiments, or with an LSM780 confocal microscope (Zeiss). WISH images were taken like Nigrosin experiments. To quantify tip GFP and mCherry fluorescence, images were first converted in 8bits, and the “threshold” function of ImageJ was used with following parameters: default, B&W, and dark background. Integrated density was then measured in a standardized area.

### Miscellaneous

Informations related to oligodeoxyribonucleotides used in this work can be found in **Table S1**. They were either synthetized by Eurofins or Sigma-Aldrich.

## DATA AVAILABILITY

Source data related to this study are available under project PRJNA1166896. The count table is available in **Dataset S1**.

## UNCROPPED BLOTS

Original files can be found in **Fig. S4**.

## ACKNOWLEDGMENTS

We thank the computing supports by the High Performance and Cloud Computing Group at the Zentrum für Datenverarbeitung of the University of Tübingen, the Baden-Württemberg state through bwHPC, and the German Research Foundation through grant no. INST 37/935-1 FUGG. The work was supported by the European Research Council under the European Union’s Horizon 2020 research and innovation programme (grant no. 757600), and intramural funding from the University of Hohenheim. Moreover, this investigation was supported by funding from the National Natural Science Foundation of China (grant no. 32388201), the Strategic Priority Research Program of the Chinese Academy of Sciences (grant no. XDB27030101), and New Cornerstone Science Foundation through XPLORER PRIZE. Finally, the work was funded by the French national research agency via grant no. ANR-20-CE2-0021, the EPIPLANT groupement de recherche (CNRS, France), and IBMP Cell Imaging Facility, member of the national infrastructure France-BioImaging supported by the French National Research Agency (grant no. ANR-10-INBS-04). We acknowledge Dr. Kerstin Feistel (Department of Zoology, University of Hohenheim) for her help in image acquisition with the LSM700 microscope, Dr. Esther Lechner (Institut de biologie moléculaire des plantes, University of Strasbourg) for having provided the mCherry antibody, and the laboratory of Dr. Laurence Drouard and Dr. Anne-Marie Duchêne (Institut de biologie moléculaire des plantes, University of Strasbourg) for helping in radioactive tRNA probe preparation. The department of Prof. Dr. Detlef Weigel (Max Planck Institute for Biology Tübingen) is acknowledged for having shared CRISPR/Cas9 vectors.

## AUTHORS CONTRIBUTION

**Conceptualization**: G.H., P.K., C.L., C.H.

**Data acquisition and analysis**: G.H., P.K., C.L., C.H., N.S., N.W., L.W., Y.X.M., E.C.K., J.M.

**Manuscript writing**: G.H.

**Manuscript editing**: All authors participated equally.

**Funding acquisition**: C.L., J.W.W., J.M.

## SUPPLEMENTARY INFORMATIONS

**Supplemental Figure 1, Sequence and secondary structure of putative mature nuclear tRNA^Tyr^ transcripts**

**A**, Alignment of mature tRNA^Tyr^ sequences. Polymorphic positions are identified by asterisks. Underrepresented NPs are highlighted with a grey background and anticodons with a yellow one. The tRNA part recognized by D/C-Tyr probes is underlined. **B and C**, Secondary structure of dispersed and clustered mature tRNA^Tyr^ species, respectively. Mismatches characterizing tRNA-like transcripts are identified with squares.

**Supplemental Figure 2, Raw deletion zone sequences**

The colour code is similar to what presented in Fig. 3A. The regions highlighted in green and yellow refer to genomic sequences in up and downstream of SYY cluster deletion, and amplified with tailed 3 + 2 primers (in red). GC/TC dinucleotides highlighted in grey in the repair zone are common to both up and downstream ends. Therefore, they origin cannot be attributed.

**Supplemental Figure 3, Consensual sequence used for the mapping of NGS data**

The colour code is similar to what presented in Fig. 3A. C-tDNA^Ser^ is highlighted in green while C-tDNA^Tyr^ ones in blue and introns in grey. Each position represented by more than one base upon SYY repeats alignment were considered as degenerated (red Ns). Y1 and Y2 positions were blocked with the MAJOR sequence for a targeted read mapping.

**Supplemental Table 1, Oligodeoxyribonucleotides used in this study**

