## Supplemental_Figures_Table for "Transcriptional reactivation of the tRNA^Ser^/tRNA^Tyr^ gene cluster in *Arabidopsis thaliana* root tip (V2)"

**Supplemental Figure 1, Sequence and secondary structure of putative mature nuclear $tRNA^{Tyr}$ transcripts**

**A**

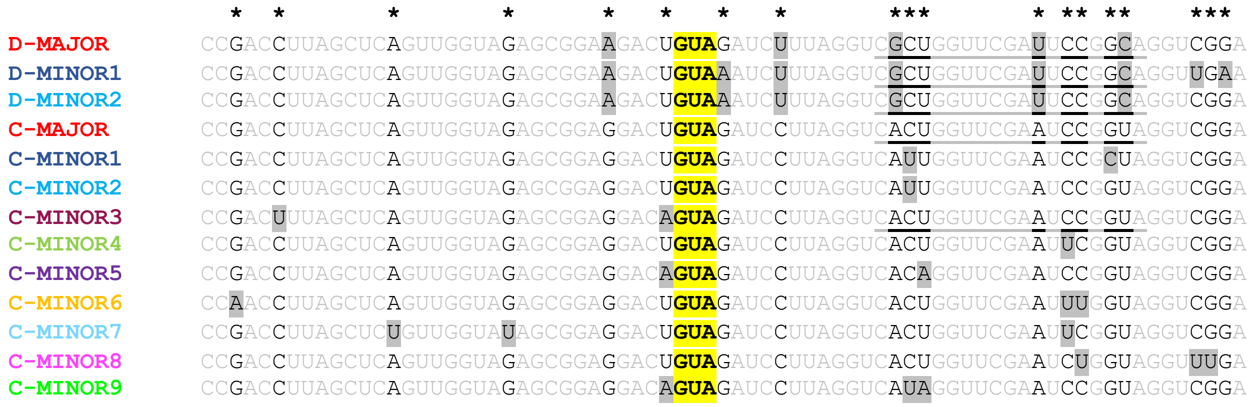


**B**

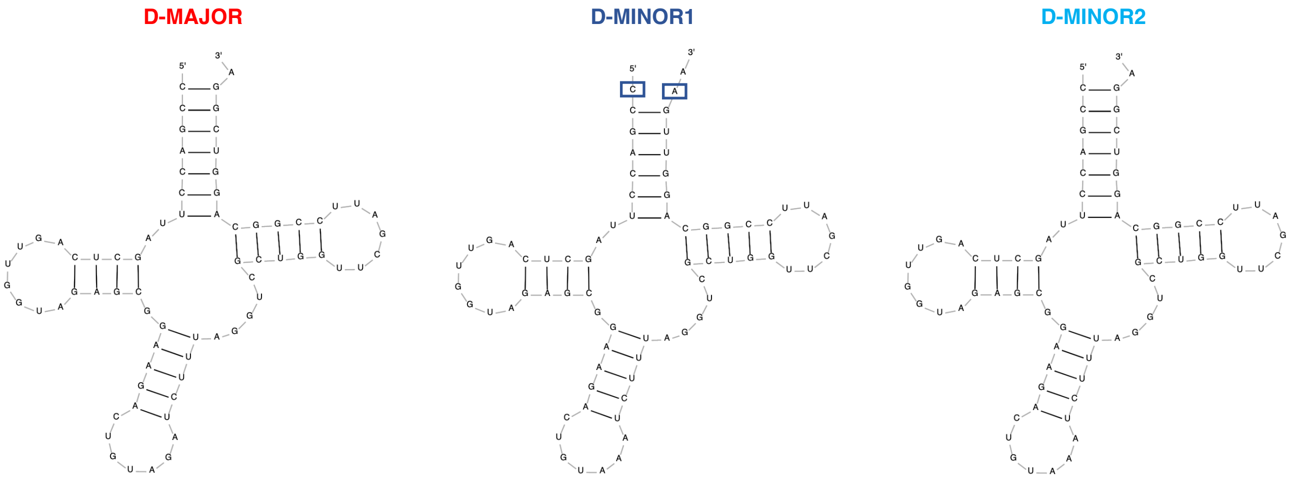

### Supplemental Figure 1, Sequence and secondary structure of putative mature nuclear tRNA$^{Tyr}$ transcripts

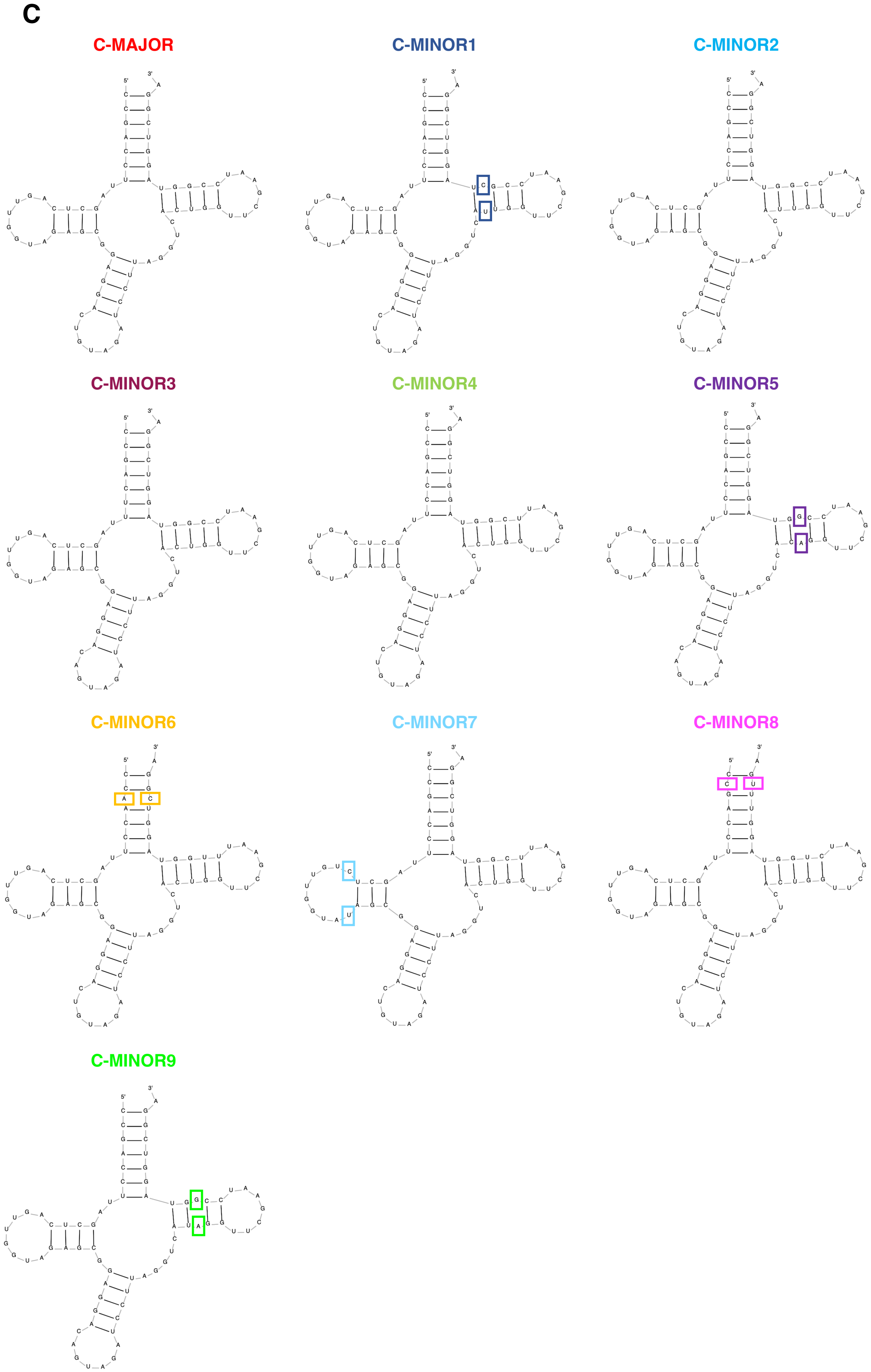

#### Supplemental Figure 2, Raw deletion zone sequences

**> T3_*syy-1*_sequencing_result_forward**

CAAACAAAGGATACGAAGAATCGTGATGTGACTGGGAGTTCCTTGGCACCCGAGAATTCCAGGTCCTGCCCTTGACACATACATATAACATA
TAGGTATTGGGCCTCTAATTTCGTGGCCCATTTTGTCTAAAATATTAATAGTGTCTTAAGAGTATCTTAAAAATAAATGAAAGAATTTAAGC
CCATTTTAGTAAAAATAAAATAGCATTTTGTTTCATTGTGAATTTTTTTTTTTTATAACCCACAATTAGCAAAATATAGAAAATTTAACAGT
TTAAAAAGCCTCCCCGAAAAATGACAAGACCGACACTGGCATCAAATTGATACATTTTCAGTTAACCGGCCATAACTCTTTAACCACAGCAC
CGGAGGATATTTAAACCGTTCGAGTTGTAGCCATCACTTTCGAATATCTACCAAATTTTTCCAAGAAGCATGTTTGCATATATTGTTGATAT
GCATTCTAATTTCTAACTTTGTTTACAATTTCTTAAAGAGGAATCACTCTGTATATTCTAAAGTACTGCAAAATCATTGTGTTCATATATTG
TTATAATTTTACACATGTATCAAATGCAATAGTACAGTTCGACCTTCGCTCAGTTGGAATTCTCGGGTGCCAAGGAACTCCAGTCACCGATG
TATCTCGTATGCCGTCTTCTGCTTGCCCTATAGTGAGTCCAAATAAAAAAA

**> T7_*syy-1*_sequencing_result_reverse_complement**

TTCACTAAAGGCAAGCAGAAGACGGCATACGAGATCGTGATGTGACTGGAGTTCCTTGGCACCCGAGAATTCCAGGTCCTGCCCTTGACACA
TACATATAACATATAGGTATTGGGCCTTTAATTTCGTGGCCCATTTTGTCTAAAATATTAATAGTGTCTTAAGAGTATCTTAAAAATAAATG
AAAGAATTTAAGCCCATTTTAGTAAAAATAAAATAGCATTTTGTTTCATTGTGAATTTTTTTTTTTTATAACCCACAATTAGCAAAATATAG
AAAATTTAACAGTTTAAAAAGCCTCCCCGAAAAATGACAAGACCGACACTGGCATCAAATTGATACATTTTCAGTTAACCGGCCATAACTCT
TTAACCACAGCACCGGAGGATATTTAAACCGTTCGAGTTGTAGCCATCACTTTCGAATATCTACCAGATTTTTCCAAGAAGCATGTTTGCAT
ATATTGTTGATATGCATTCTAATTTCTAACTTTGTTTACAATTTCTTAAAGAGGAATCACTCTGTATATTCTAAAGTACTGCAAAATCATTG
TGTTCATATATTGTTATAATTTTACACATGTATCAAATGCAATAGTACAGTTCGACCTTCGCTCAGTTGGAATTCTCGGGTGCCCAAGGAAC
TCCCAGTCACCCGATTATCTCGTAGCTTTTCGGC

**> T3_*syy-2*_sequencing_result_forward**

CACCAAAGGAACGAAGAATCGTGATGTGACTGGGAGTTCCTTGGCACCCGAGAATTCCAGGTCCTGCCCTTGACACATACATATAACATATA
GGTATTGGGCCTCTAATTTCGTGGCCCATTTTGTCTAAAATATTAATAGTGTCTTAAGAGTATCTTAAAAATAAATGAAAGAATTTAAGCCC
ATTTTAGTAAAAATAAAATAGCATTTTGTTTCATTGTGAATTTTTTTTTTTTATAACCCACAATTAGCAAAATATAAAAAATTTAACAGTTT
AAAAAGCCTCCCCGAAAAATGACAAGACCGACACTGGCATCAAATTGATACATTTTCAGTTAACCGGCCATAACTCTTTAACCACAGCTCGG
ATTAACCCACAATGACCTACAAAAACATACAAAACACACGAAAAGACACACAAAAACACTATAGAACTTCTAAAATGGAAAAAAACTCACTT
GGACTCTAAATAGACAAAACATAAGGAAAATACCAATAAAATACGATAAAAAATGATATTAAAAACCGTGATAAAGCGGTAAATATAGAGTA
TATCAGCAAGTTCGACACATGCCACTATGCCAGCTTACTAAAATATGGCGTCAAGGGTTTGTACAAGACTAATAAGCACGTCAATGCCTGAT
AGCTCGTAAGATATAGAAAGTAAAACACCTACAAATGCCTTTTCGTAAGAAGGTGTAAGAAATCTAGTGGAAACACTAAAAGAGGCAACCCT
GGTTCTCGGAAAACAGACGCAAGATCCTTCATGTTTGTTTATCGTGACCAAGATGTATTTTGAAGATTTCCAATCAGGCCCGGCCCAAGTGG
GTAGGGTAGAGGTGCTTAGGTTTAAAGCTCATCGAGTTGTAGCCATCACTTTCGAATATCTACCAGATTTTTCCAAGAAGCATGTTTGCATA
TATTGTTGATATGCATTCTAATTTCTAACTTTGTTTACAATTTCTTAAAGAGGAATCACTCTGTATATTCTAAAGTACTGCAAAATCATTGT
GTTCATATATTGTTATAATTTTACACATGTATCAAATGCAATAGTACAGTTCGACCTTCGCTCAGTTGGTAGACTAGGAGGTTGTTTCTTTG
CGCCAAGAACTGTGCAAAAACCACATGTGGGAGGGATCTTTTGGGTTTTGGAAGCCGGCCTAGTCAAGGGTTCAAACATCAAAAGGGGGCAA
GTAAAGGGAAAGTAACCTCTGAAATTTCCCCCCATTCCCTATAAAAAATTGGGAGGGAAGGGACTTTACCTTTCCAAAAAAAAAACTTGGAT
TTGGTTTTCCTTTTGGTTCCTTTTTAAAAAAATTTTTGGAAACCTTTGCAAAAAACCGAAAAGGGAAGAAAAACATTTTTAAAAAAAAAAAT
CACCCCCCCCGGATAAAAAATTTTCCCCAGGGGTGGGGTTTCCTTATGTTTTTTTTAAAAAAAAAAAAAACCCTTC

**> T7_*syy-2*_sequencing_result_reverse_complement**

TTTTTTATAACCCAAAAAAGGCAGCCCCTCTATTTATTTTTTTTTTTGCCCCGCCTTTTTTTCCCCGGGTTGGTTCCCCCTAGTTTTTAAAA
AAATTAACACCCGAAAACCCCAAAAACCCTTTATTTTTTTTAAAAAAGAAAAACATCCCCCTTGTGTCTCTAAAACACAAAACCTGGGGAAA
CCCCCCAAAAATGGGTAAAAAATTGTTTTAAACCCGGGAAAAAGGCGGTAATTTGAGGTTAATCCCCATTTTCGCCCCCTCGCCCCTTTTCC
CCGCTTTTTTAAAAGGGGTCAAGGGTTTTTACAAAATTAAAAAGCCCGTCAATCCCTGATAGTTTGTAAGATTAGAAAGTAAAACCCCTACA
AATGCCTTTTTGTAAGAAGGTGTAAGAAATCTAGTGGAAACACTAAAAGAGGCAACCCTGGTTCTCGGAAAACAGACGCAAGATCCTTCATG
TTTGTTTATCGTGACCAAGATGTATTTTGAAGATTTCCAATCAGGCCCGGCCCAAGTGGGTAGGGTAGAGGTGCTTAGGTTTAAAGCTCATC
GAGTTGTAGCCATCACTTTCGAATATCTACCAGATTTTTCCAAGAAGCATGTTTGCATATATTGTTGATATGCATTCTAATTTCTAACTTTG
TTTACAATTTCTTAAAGAGGAATCACTCTGTATATTCTAAAGTACTGCAAAATCATTGTGTTCATATATTGTTATAATTTTACACATGTATC
AAATGCAATAGTACAGTTCGACCTTCGCTCAGTTGGTAGACTAGGAGGTTGTTTCTTTGCGCCAAGAACTGTGCAAAAACCACATGTGGATG
GATCTTTTGGTTTTGTAGTCTGCCTAGTCAATGGTTCAAACATCATAAGGTGTCAAGTATAGTGAAAGTAACCTCTGACATTTCCACTCATT
CCCTATAACAAACTTGGCATGTAACGTAACTTAACCTTTTACCAAAAAAAAAAAACTTGACATGTATGCATTTCTCTTATTGAGTTCTCTTT
TGATAACGATTTTGTTGTAATACTCTTGTCAAAGAACACTGAAAATGGAATGAAACAAGCTTTAGAAAATAAGATCTGACCCTACCGGATTC
AAACTATTGACCTAAGGATGTTCTGCTTCAACTATTGTTTTCATTCGAAATGCAAAGAAACAACCTCCGTTCCAAAAAGCTAGATCTTTGGC
CTCCATAGCTTGCAGAATTCACTTGATTAAGTTATGTTCTCCATGACATTATAATACCACAGGCTTATATGCTGTGTCTATATGACAATATC
ACTGAAATCCTAGCTCTTTACTCACATTCAAAACACATTGATGTCGATTATAATTAATTAATTAATTATGCTCGAGAAATGGTAGCATTAGG
TTTGTTGGTGATCAAACATATTCCTACACATATTTAACTAGCAGGCATCTCCATGAAGTCACAACCTCGGTGTGGTGGAATTCTCGGGTGCC
AAGGAACTCCCAGTCACCCGATGTATCTTCGTTGCTTTCTGT

#### Supplemental Figure 3, Consensual sequence used for the mapping of NGS data

CNAGGGTNGTGAAATANCNTTTNNNACTNTGTTGCNNGNATTTGTTTGGCTTTATTTTCAAGAATATAAACAATNATNNGCNCANTCTCAATATTACATGCATAAATATANCATCAAAGTTGCTCTTTA
GAAATAATTTANTTTTTGANNTGAGAAAATNTTATAAAAATTNNAAANAATCAAGACAGCCAAATAGGTTTATGGTTCTNATNTTTTTTATTNTGTTCATAAACTNNNTNATCCTTGNAATNTTCTACT
TCTNACTCATGATCAAAATTTAGGATANAGAATGGTATANNGTTTAGTCTATTTTGAAATAAAAGTTTAGTTTANAGAAAANGTGGANNTGNCGGAGTGNNGTTATCGGGNATNACTAGAAATCATGNN
GGNTTTGCCCGCGCANGTTNGAATCNTGCCGTTNACGTTTTTATTTTGAGTGTTGGTNAGTTNTTANAATNACTTANTGTGTTTTATGNATNGTNTTNNAATTTGAACNGTTTTNCATCATGTTACNGT
TTTCAGTAAANTTTAGTATTGATNAAAAAANNTNNNNAAAAAACAANANAGCATGAAAGATATGAAGTCAACTTTNTTTCTANTCCTCTTCNNNANAAGTTTAAATTTGATTAAAAAGNTNNNNTNGAA
TCATCAACATGCNTAAAGTGTTATNATACNAAACCGACCTTAGCTCAGTTGGTAGAGCGGAGGACTGTAGTTGANGCAGATNATCCTTAGGTCACTGGTTCGAATCCGGTAGGTCGGATCATNAAANTT
NAAANNNTTNTTTNNTNCATTTCGTTTCAAAGNGNTTAGANAAGAGTATAACTACAAACTNNTTTTTCTATNANAANTTNGTTTNAGAGAATTGCATAGCTANTGANNGTATTATCANAAANGAGTGGG
AANTCTNNAAGNAATTTTCACTGTANTTTAAACCGTTNAAGNNNNNNNNNNNNNNNNNNNNNNNTTACGAGNTTTANGCAANNNNNNNNGNNTTTTTCNAATATAATGTTTTNATAAATTCAAACTTTN
TTTNCAAANTTNTAATANAGANTCACTATGANTATGCTAACTTAATACAAATCATTNTGNTNATAGAATATTTNGATCAGTACACATGCATGAAATANAATACAATCCGACCTTAGCTCAGTTGGTAGA
GCGGAGGACTGTAGNAGACGNAGATTATCCTTAGGTCACTGGTTCGAATCCGGTAGGTCGGAATTTGCTCCCACANGAGANCTTTTTATTTTTCTTTNGNTGTGACATTAANNTNTTTNNAATTTTATA
ATAAACGGTTATATGGTGGTCGACAATTNAACATACCNAAAAGTTTNGCTCGNATTNTTNAANGATCCGTATGTTTANCNTTTTTCAATATTGATACGATGANAAATGATTTAAAAGTGANTAAAAACT
ATGAGTTTTCTAATTTNCTTTGGTCAAACAAANATTTAGTTTTANAGTTTTAAAAANNACAAANAATGTNTGCATANTTATNTTCAAATGCTTGTGGGTTGTGCCTATAAGTTGTCAACGTTCATA

**Supplemental Figure 4, Uncropped blots**

**Figure 2B,** Roti staining

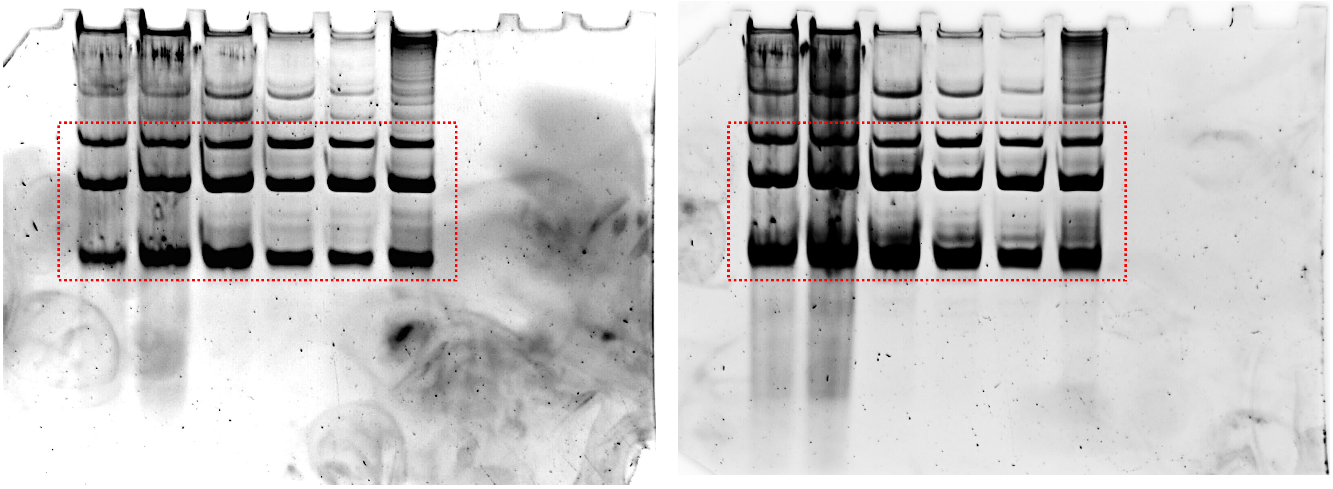

**Figure 2B,** D/C-Tyr E1

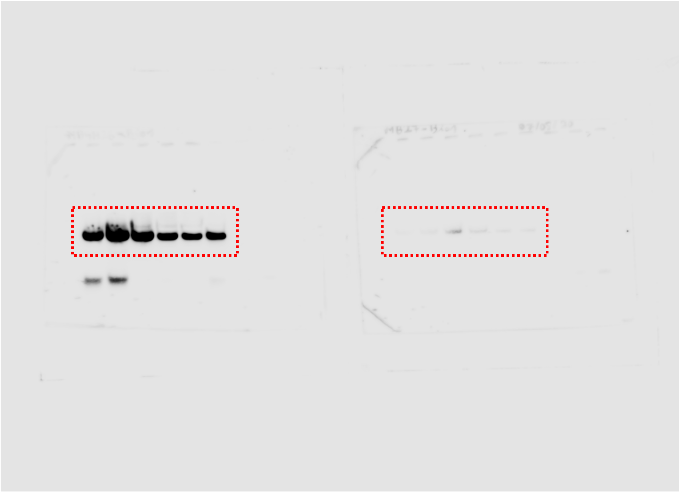

**Figure 2B,** D/C-Tyr E2

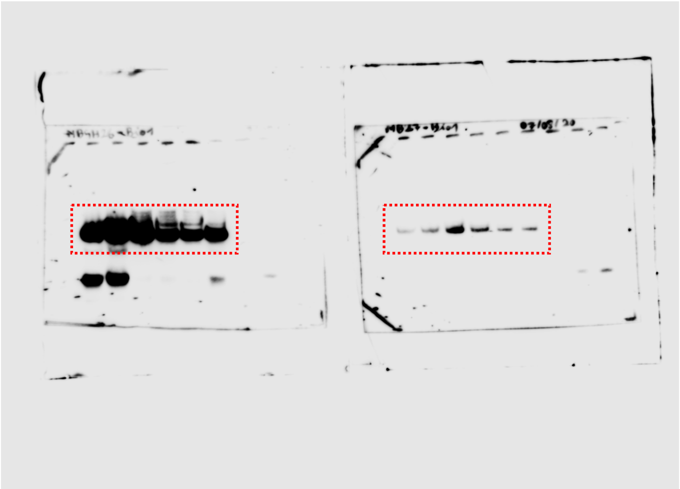

**Figure 2B,** D-Ala

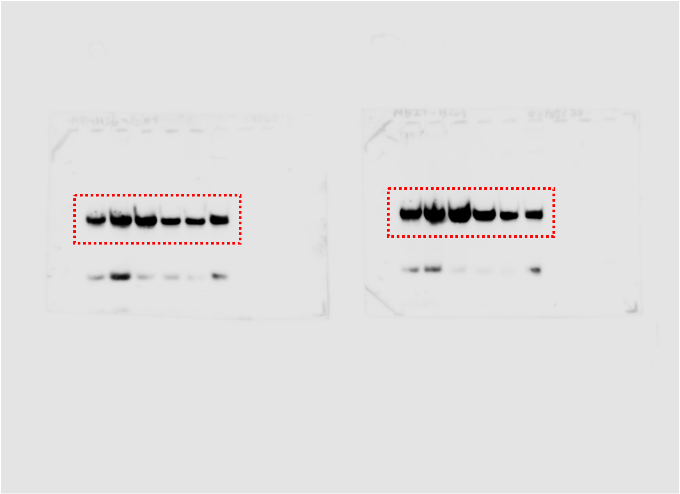

**Supplemental Figure 4, Uncropped blots**

**Figure 2D,** Roti staining

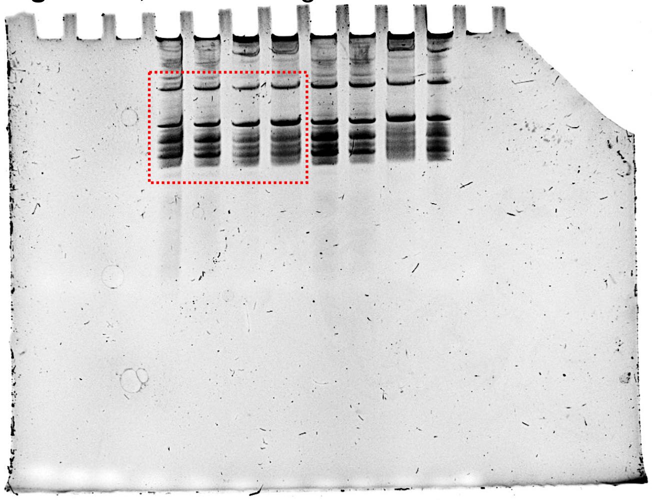

**Figure 2D,** C-Tyr

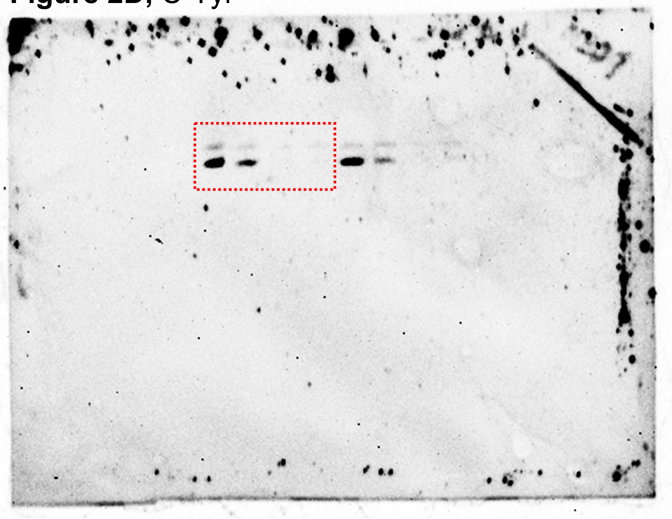

**Figure 2D,** D-Ala

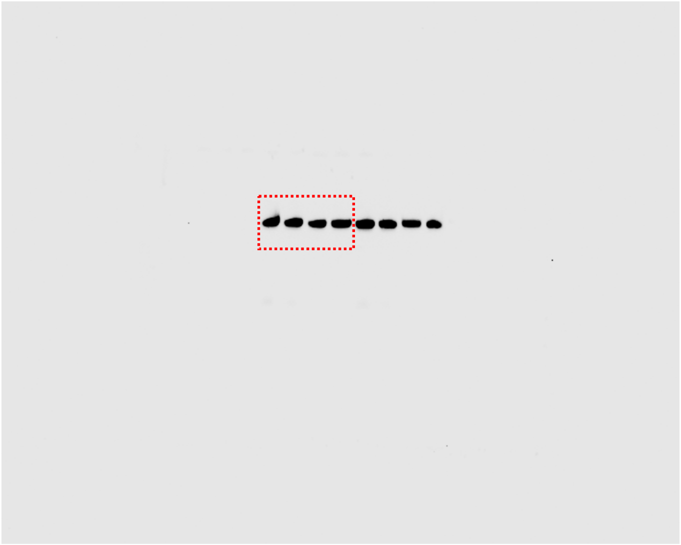

**Supplemental Figure 4, Uncropped blots**

**Figure 3C,** Roti staining

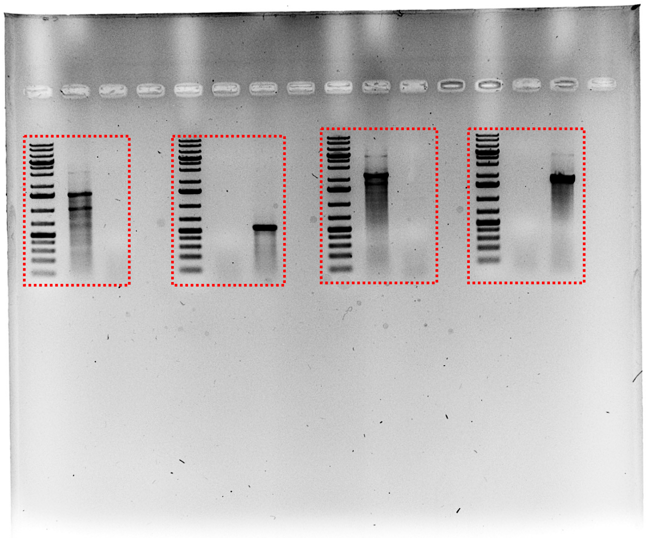

**Figure 3D,** Roti staining

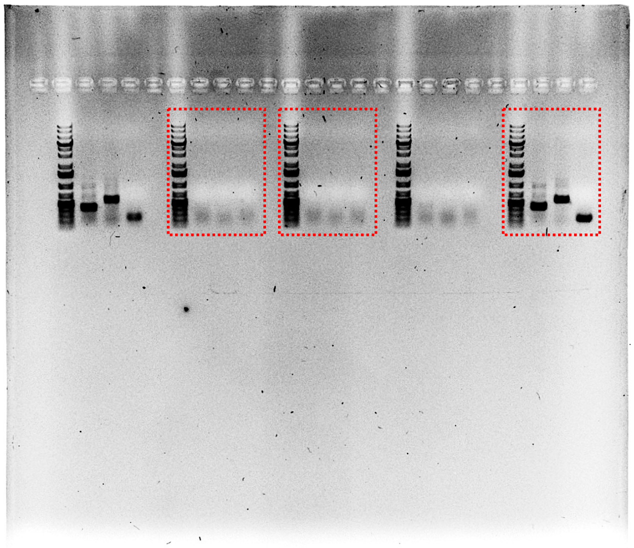

**Supplemental Figure 4, Uncropped blots**

**Figure 3G,** Roti staining

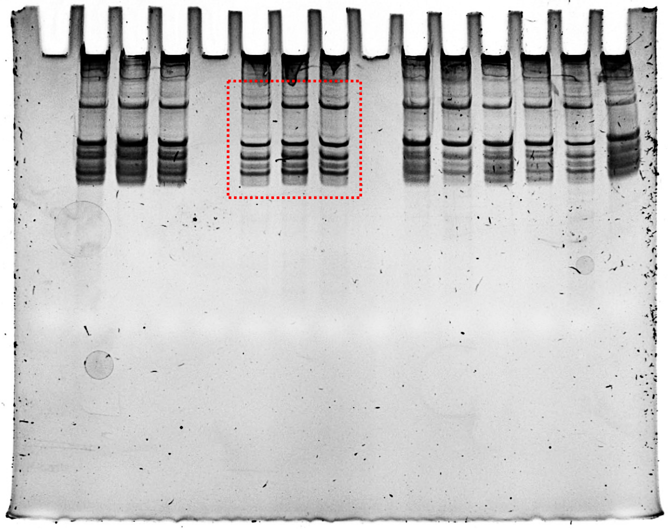

**Figure 3G,** D-Tyr

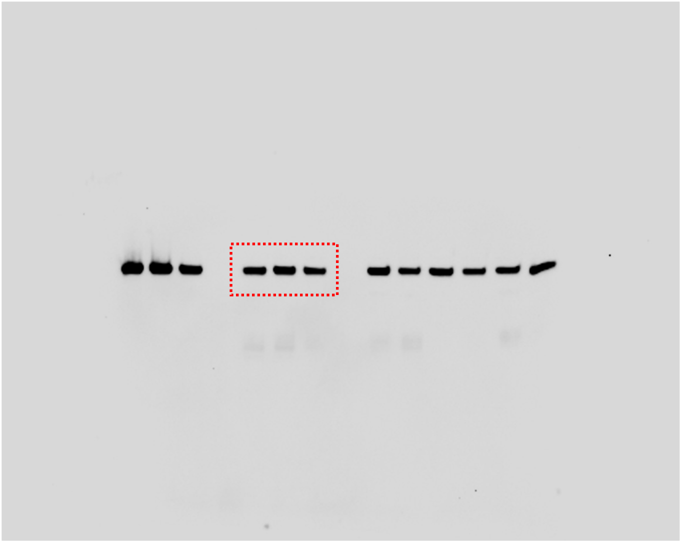

**Figure 3G,** C-Tyr

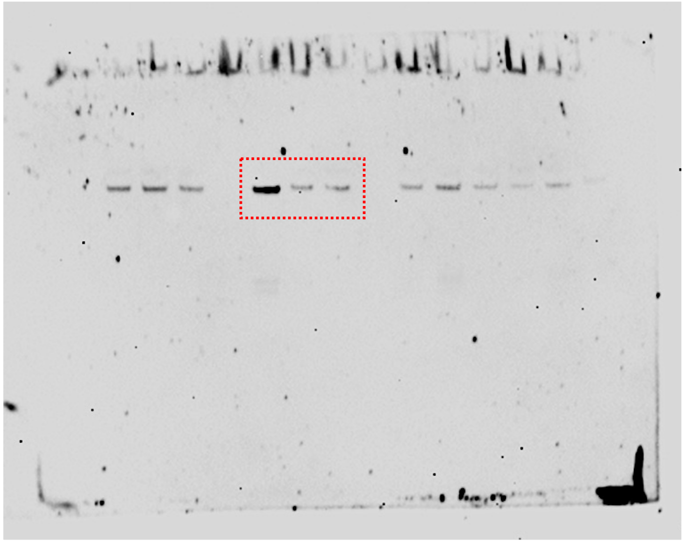

**Figure 3G,** D-Ala

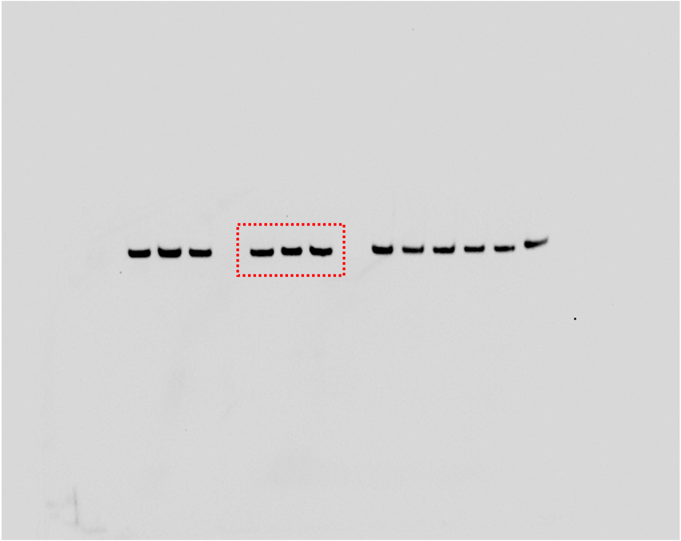

**Supplemental Figure 4, Uncropped blots**

**Figure 5E,** p probe

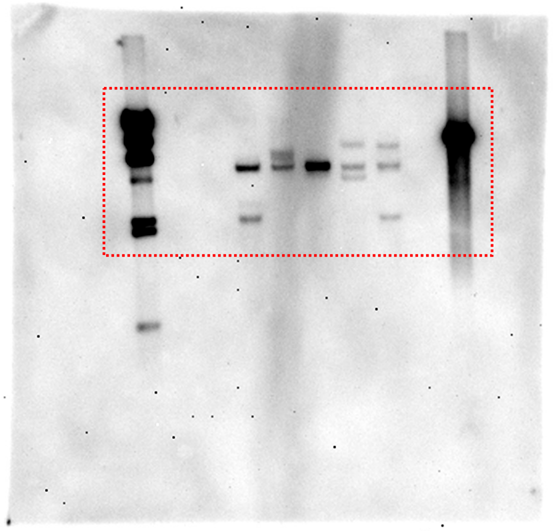

**Supplemental Figure 4, Uncropped blots**

**Figure 5E,** Roti staining

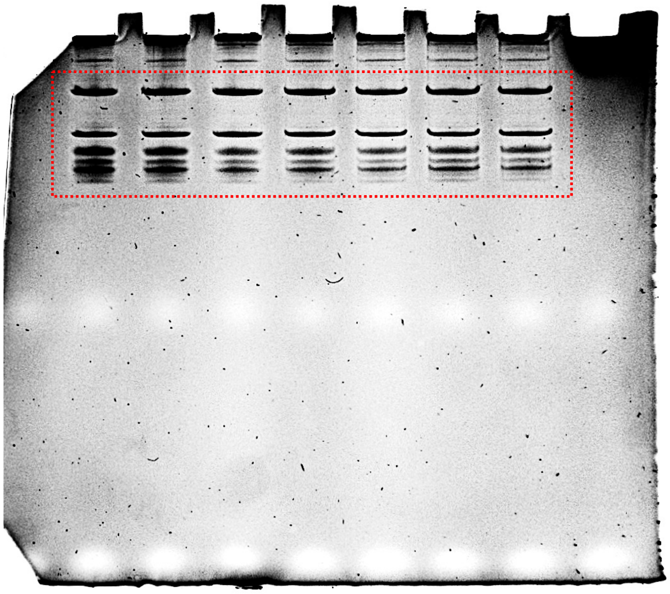

**Figure 5E,** C-Tyr

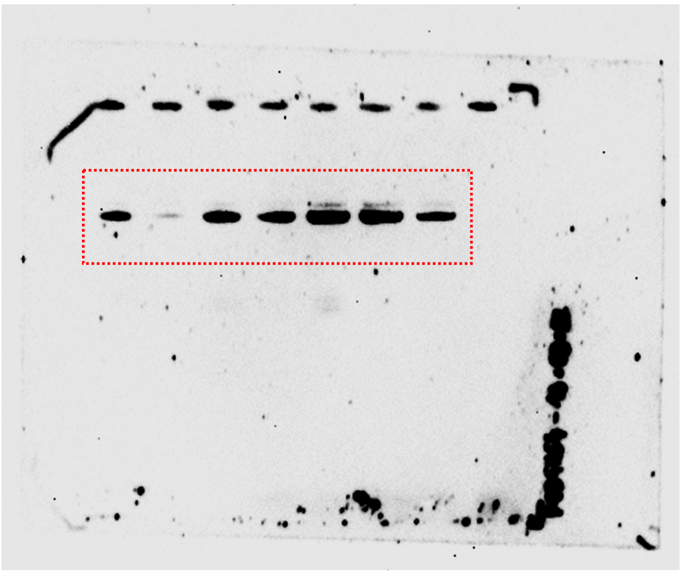

**Figure 5E,** D-Ala

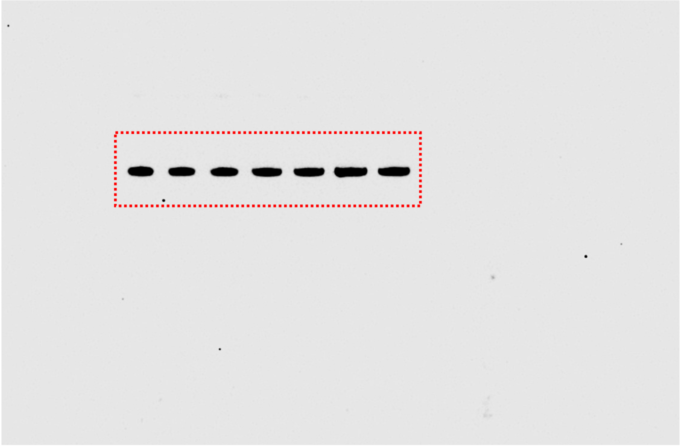

**Supplemental Figure 4, Uncropped blots**

**Figure 5G,** Roti staining

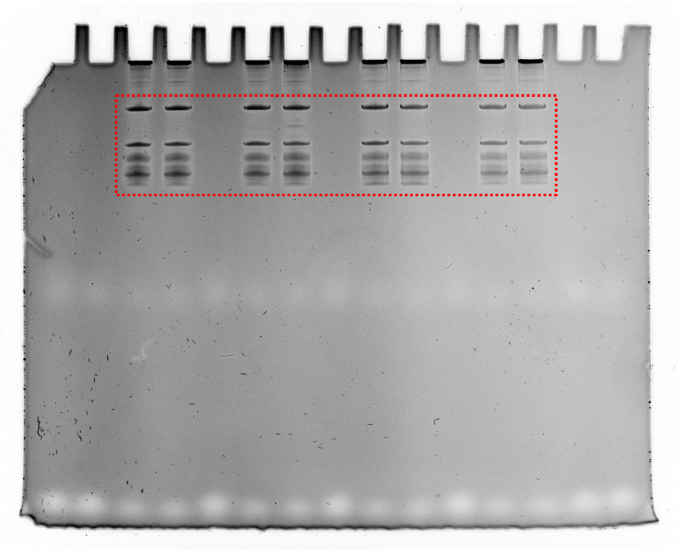

**Figure 5G,** C-Tyr

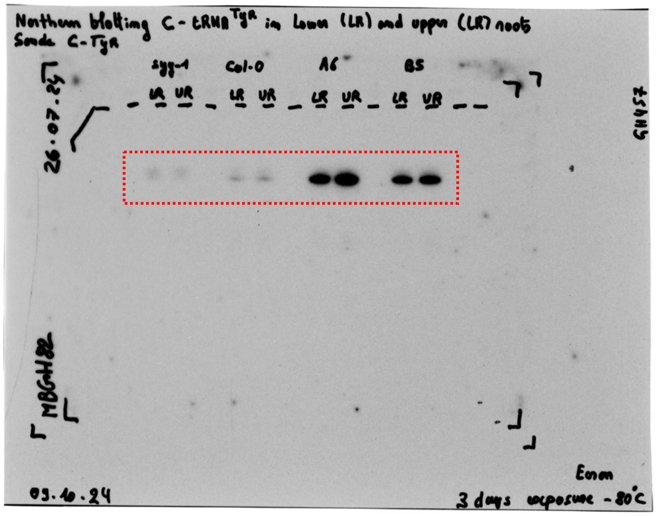


**Figure 5G,** D-Ala

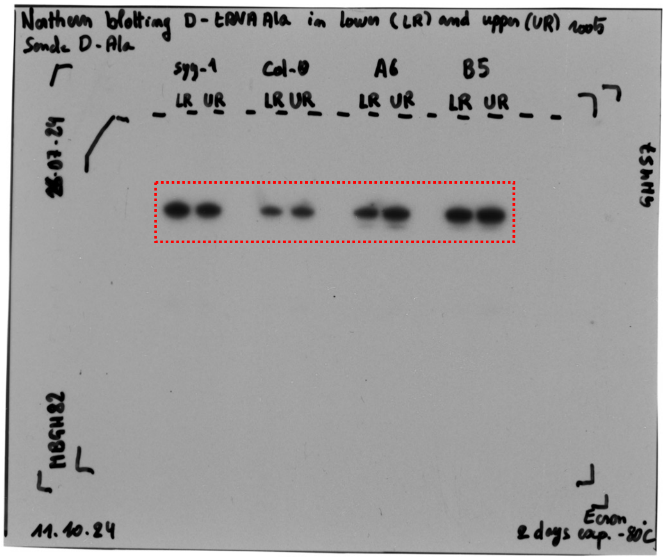

**Supplemental Figure 4, Uncropped blots**

**Figure 6C,** Roti staining Val PCR

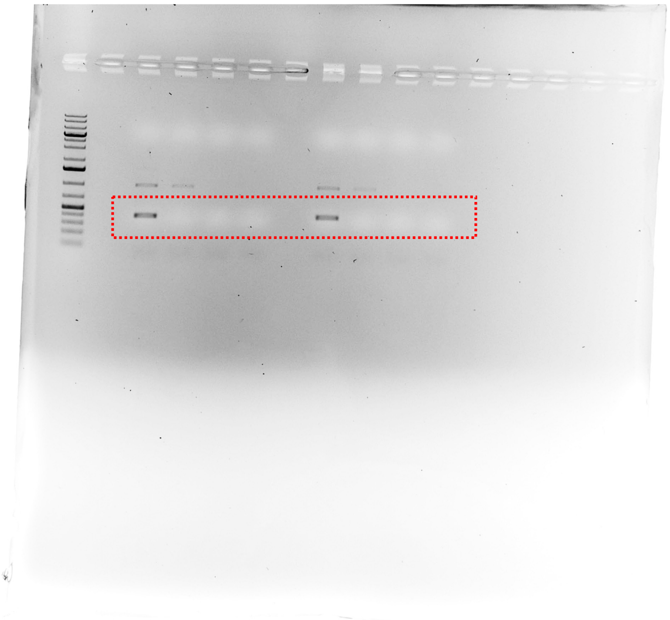

**Figure 6C,** Roti staining Y1 PCR

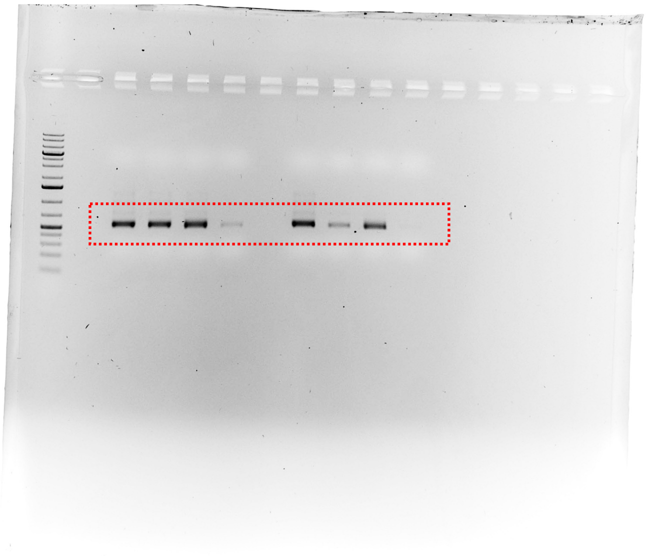

**Figure 6C,** Roti staining Y2 PCR

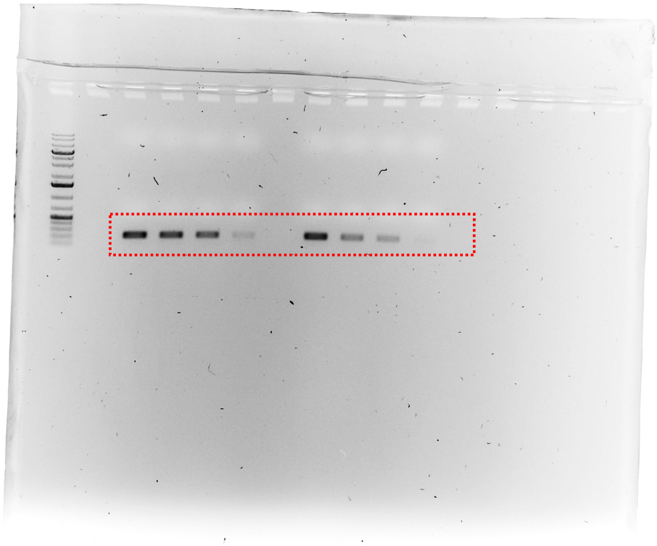

**Supplemental Figure 4, Uncropped blots**

**Figure 6H,** Roti

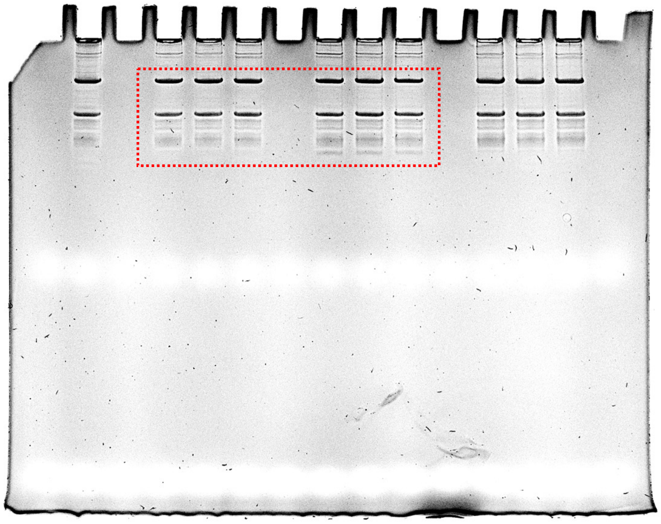

**Figure 6H,** C-Tyr

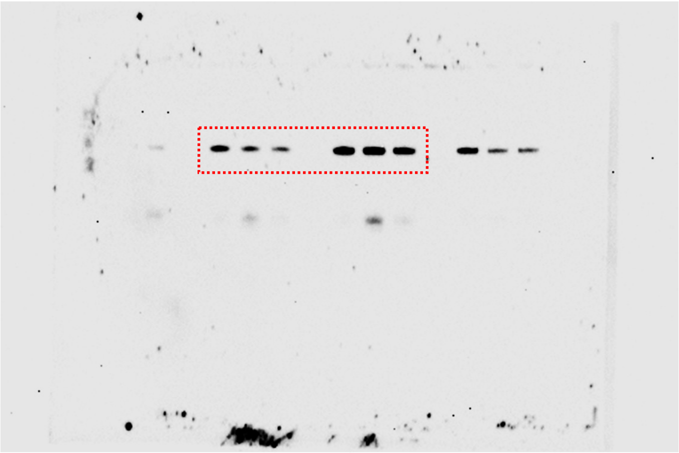

**Figure 6H,** D-Ala

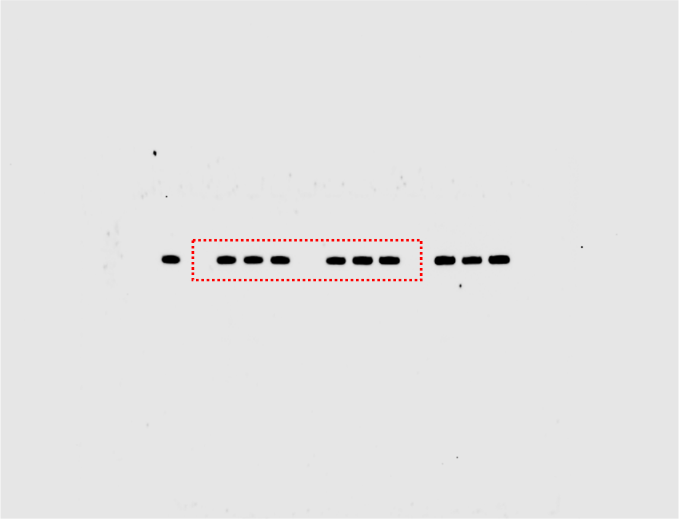

**Supplemental Figure 4, Uncropped blots**

**Figure 7A,** Roti staining Val PCR

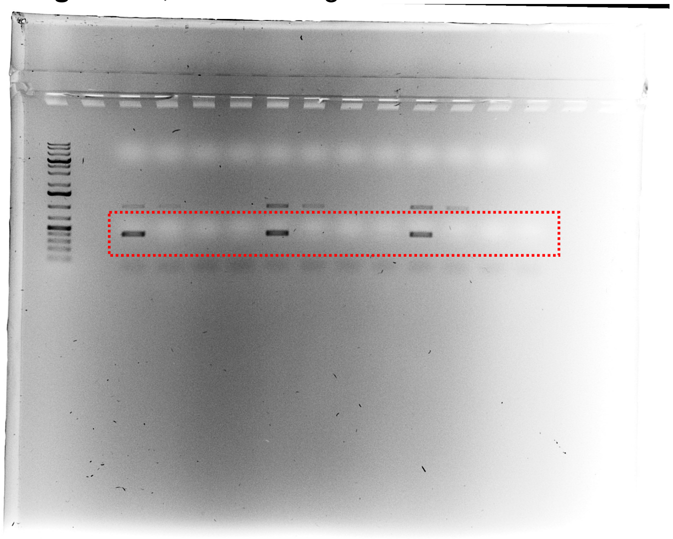

**Figure 7A,** Roti staining Y1 PCR

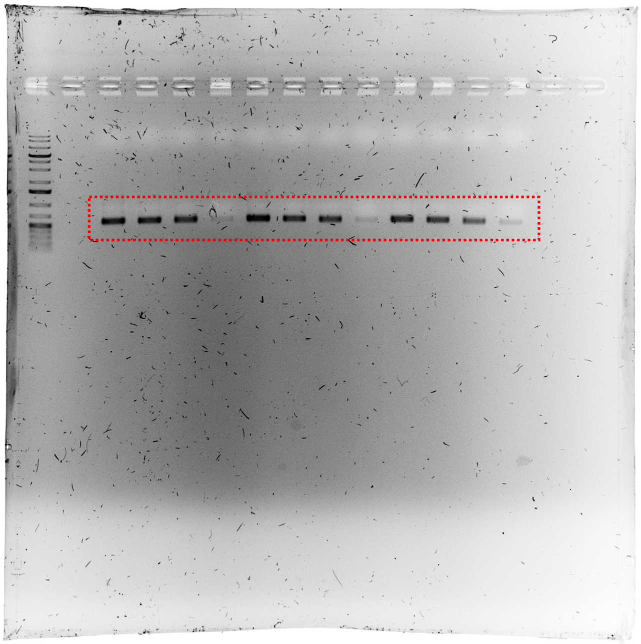

**Figure 7A,** Roti staining Y2 PCR

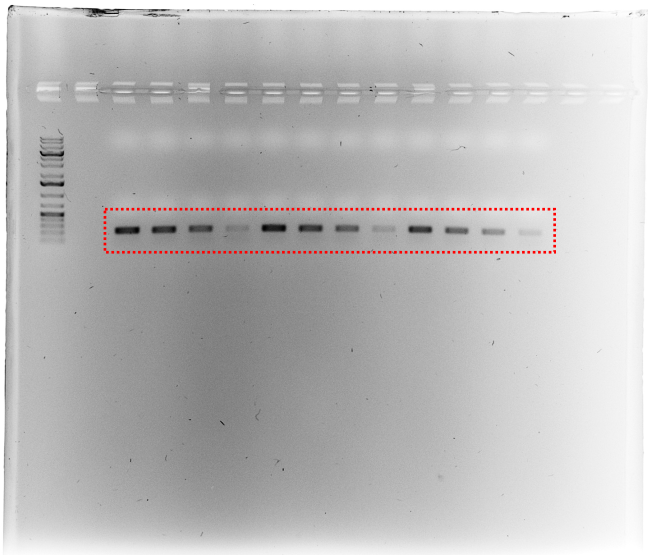

**Supplemental Figure 4, Uncropped blots**

**Figure 7C,** Roti staining Val PCR

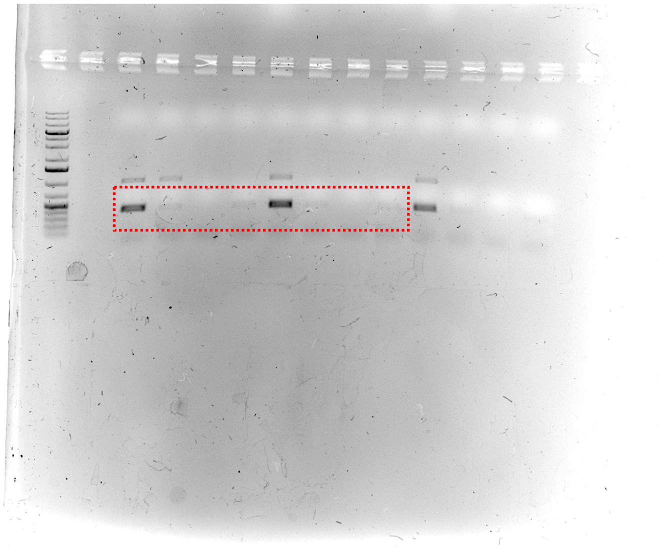

**Figure 7C,** Roti staining Y2 PCR

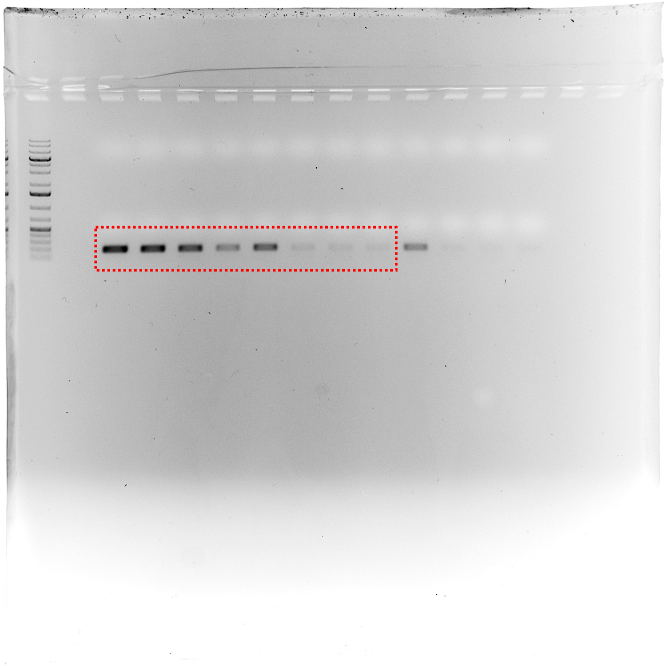

**Supplemental Figure 4, Uncropped blots**

**Figure 7F,** Roti staining

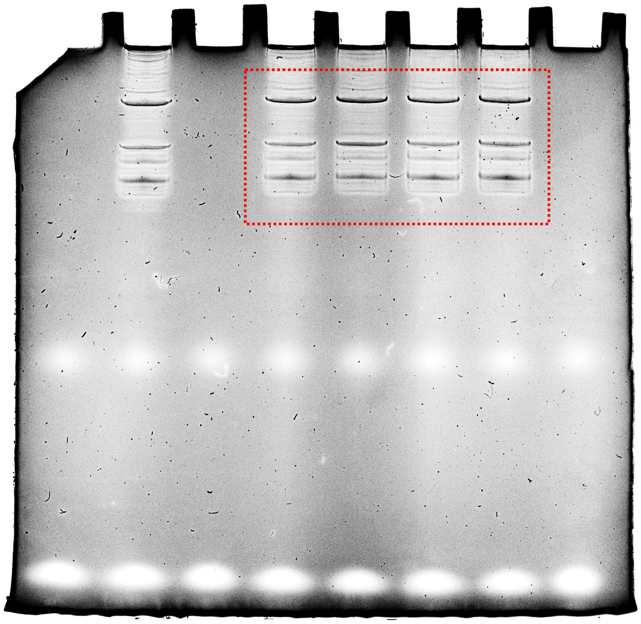

**Figure 7F,** C-Tyr

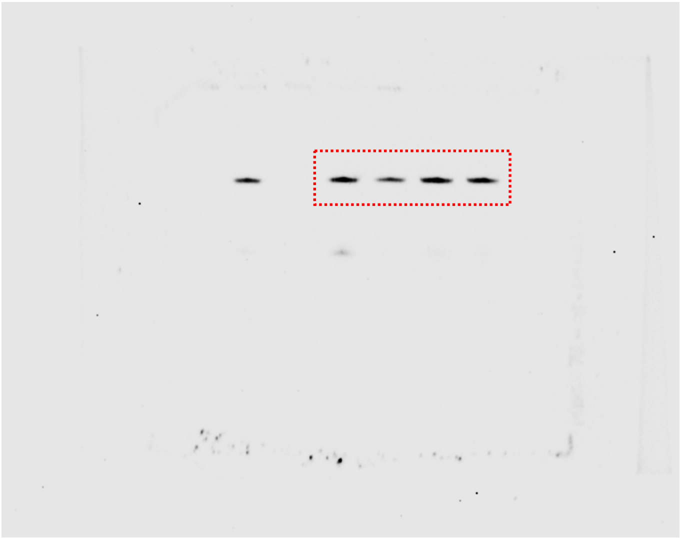

**Figure 7F,** D-Ala

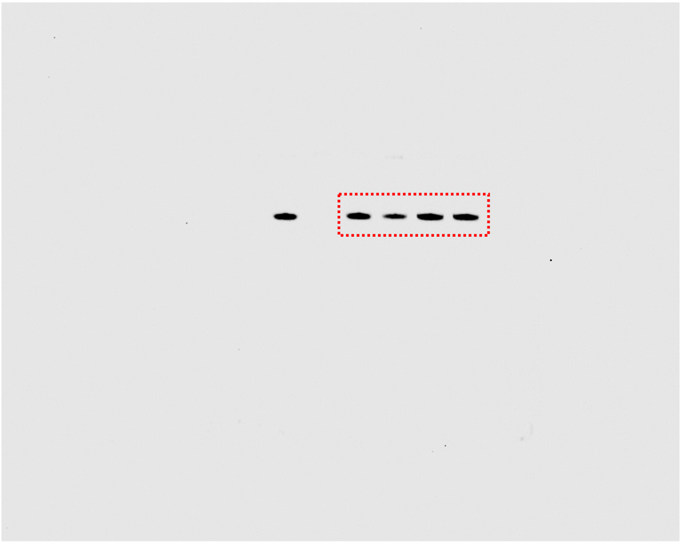

**Supplemental Figure 4, Uncropped blots**

**Figure 10F,** Coomassie staining

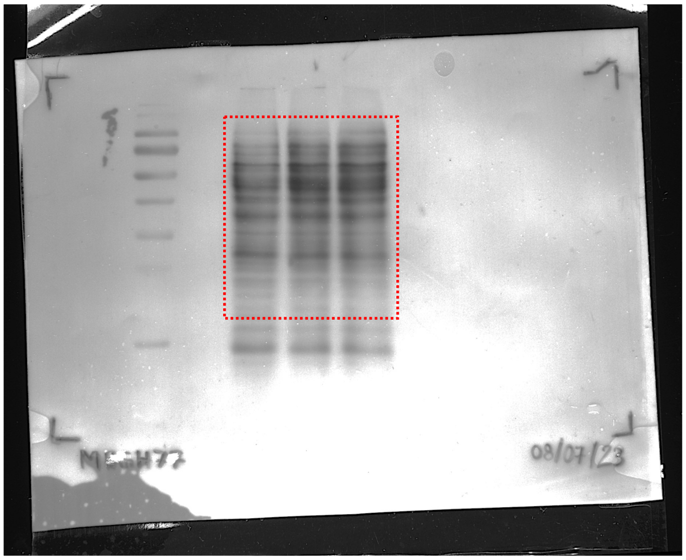

**Figure 10G,** Coomassie staining

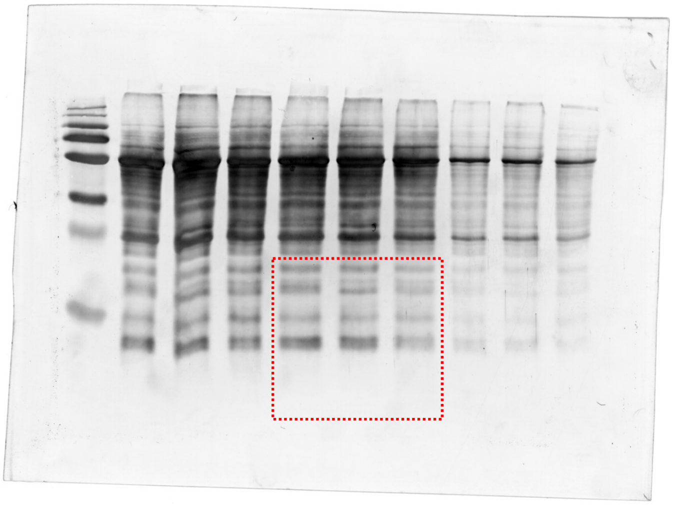

**Figure 10F,** Western blot @GFP

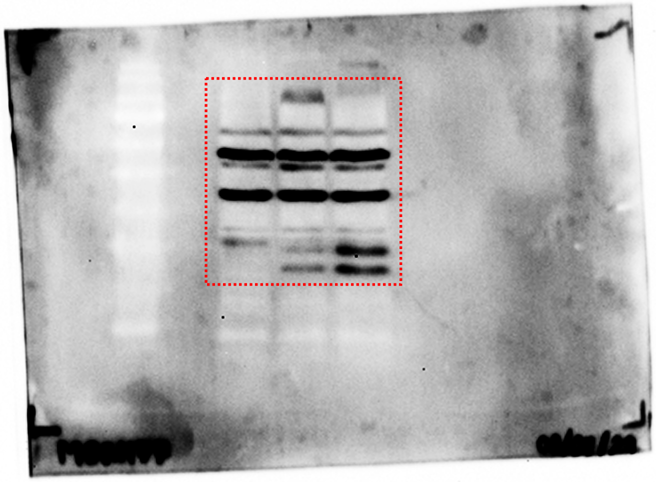

**Figure 10G,** Western blot @mCherry

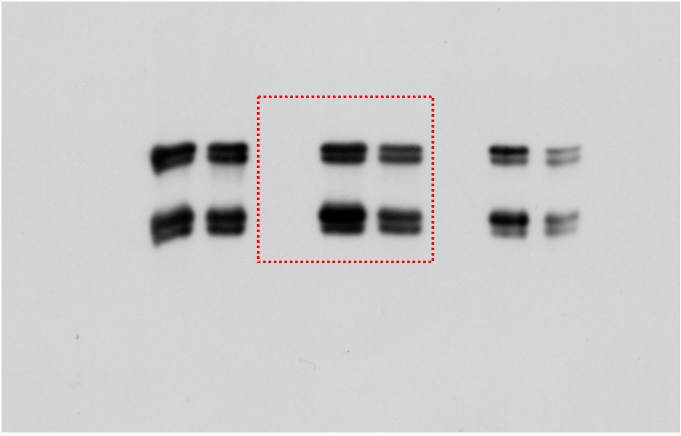

#### Supplemental Table 1, Oligodeoxyribonucleotides used in this study

| Aim(s) | Name | Sequence (5'-3') and specifications |
|---|---|---|
| CL | GH1-sgRNA | TTGTGGGTTAATCCGAGCTG |
| CL | GH2-sgRNA | AACTGGTAGGCCTTGGGCAC |
| CL | Fw-pEF004-GH1-sgRNA | TTGTGGGTTAATCCGAGCTGGTTTTAGAGCTATGCTGAAAAG |
| CL | Rv-pEF004-GH1-sgRNA | CAGCTCGGATTAACCCACAACAATCACTACTTCGACTCTAGC |
| CL | Fw-pEF005-GH2-sgRNA | AACTGGTAGGCCTTGGGCACGTTTTAGAGCTATGCTGAAAAG |
| CL | Rv-pEF005-GH2-sgRNA | GTGCCCAAGGCCTACCAGTTCAATCACTACTTCGACTCTAGC |
| CL | LW1-sgRNA | TCGAGGAAACAAATCCCAA |
| CL | LW2-sgRNA | TATATTTACCGCTTTATCA |
| CL | LW3-sgRNA | GGTTTAAAGCTCATCAATA |
| CL | LW4-sgRNA | CAATCATCGACCATTTGGG |
| CL | LW5-sgRNA | GTGATGGCTACAACTCGAA |
| CL | LW6-sgRNA | CGCTCAGTTGGTAGACAGG |
| CL | Fw-atU3b-LW1-sgRNA | TCACGTCGAGGAAACAAATCCCAA |
| CL | Rv-atU3b-LW1-sgRNA | AAACTTGGGATTTGTTTCCTCGAC |
| CL | Fw-at7SL-LW2-sgRNA | TTACGTATATTTACCGCTTTATCA |
| CL | Rv-at7SL-LW2-sgRNA | AAACGTGATAAAGCGGTAAATATAC |
| CL | Fw-atU6-LW3-sgRNA | GATTGGGTTTAAAGCTCATCAATA |
| CL | Rv-atU6-LW3-sgRNA | AAACTATTGATGAGCTTTAAACCC |
| CL | Fw-atU3b-LW4-sgRNA | TCACGCAATCATCGACCATTTGGG |
| CL | Rv-atU3b-LW4-sgRNA | AAACCCCAAATGGTCGATGATTGC |
| CL | Fw-at7SL-LW5-sgRNA | TTACGGTGATGGCTACAACTCGAA |
| CL | Rv-at7SL-LW5-sgRNA | AAACTTCGAGTTGTAGCCATCACC |
| CL | Fw-atU6-LW6-sgRNA | GATTGCGCTCAGTTGGTAGACAGG |
| CL | Rv-atU6-LW6-sgRNA | AAACCCTGTCTACCAACTGAGCGC |
| CL | Fw-pFK206-attB2-SV40NLS | CCGAAGAAGAAGCGAAAGGTCTAAACCCAGCTTTCTTGTACAAAG |
| CL | Rv-pFK206-attB1 | GAAAGCCTGCTTTTTTGTACAAACTTGTGATCCTCTAGAACTAGTGGATC |
| CL | Fw-attB1-pUBQ10-5'UTR | GTACAAAAAAGCAGGCTTTCAGTCTAGCTCAACAGAGCTTTTAACC |
| CL | Rv-pUBQ10-5'UTR | CTGTTAATCAGAAAAACTCAGATTAATCGACAAATTCGATCGCACAAACTAGAAACTAACAC |
| CL | Fw-pUBQ10-5'UTR-EXT19CDS | TGAGTTTTTCTGATTAACAGATGCAATACACATACTCTCCACCATCAC |
| CL | Rv-EXT19CDS-2xGly | TCCGTATATTGGAGGAGGGGGTGAGCTATAGCTATAGC |
| CL | Fw-EXT19CDS-2xGly-GFPCDS | CCTCCAATATACGGAGGAATGAGCAAGGGCGAGGAGC |
| CL | Rv-GFPCDS-SV40NLS-STOP | CCTTTCGCTTCTTCTTCGGCTTGTACAGCTCGTCCATGC |
| CL | Fw-pUBQ10-5'UTR-mCherryCDS | TCGATTAATCTGAGTTTTTCTGATTAACAGATGGCCATCATCAAGGAGTTCATGCGCTTC |
| CL | Rv-mCherryCDS-SV40NLS-STOP | ACCTTTCGCTTCTTCTTCGGCTTGTACAGCTCGTCCATGC |
| CL | Fw-attB2-pFK206-p2S3-mCherry | ACCCAGCTTTCTTGTACAAAGTGGTTCGGAGCTCCAGCTTTTGTTC |
| CL | Fw-attB1-Ing-Y | ACAAGTTTGTACAAAAAAGCAGGCTTTCATTGGTAAGTGAATGGTTGTAG |
| CL | Rv-attB2-Ing-Y | ACCACTTTGTACAAGAAAGCTGGGTTTTTCAACCATTCAAGTGAGAGAG |
| CL | Fw-attB1-1xSYY | ACAAGTTTGTACAAAAAAGCAGGCTTTCCCCAAGTGGGTAGGGTAGAG |
| CL | Rv-attB2-1xSYY | ACCACTTTGTACAAGAAAGCTGGGTCGTGCAACAGAGTGGGAAAG |
| GN | Fw-1 | GGACTGTAGTTGACGCATCCTTAG |
| GN | Rv-2-*syy-1* | ACTGAGCGAAGGTCGAACTG |
| GN | Rv-2-*syy-2* | CCACACCGAGGTTGTGACTTC |
| GN | Fw-3 | GGTCCTGCCCTTGACACATAC |
| CP + GN | Fw-α | GACAGCCAAATAGGTTTATGGTTC |
| CP + GN | Rv-β | AAAGTTGACTTCATATCTTTCATGC |

| | | |
|---|---|---|
| CP + GN | Fw-γ | AGCATGAAAGATATGAAGTCAAC |
| CP + GN | Rv-δ | TATTTCATGCATGTGTACTGATC |
| CP + GN | Fw-ε | GATCAGTACACATGCATGAAATA |
| CP + GN | Rv-ζ | AATTGTCGACCACCATATAACC |
| CP | Fw-Ala | AACATTTCGACGCGAGAGC |
| CP | Rv-Ala | TGCCGTTGGTTTGAAGTTGG |
| CP | Fw-Val | GTTGTCCAAGTTCCGTCTAC |
| CP | Rv-Val | GATGATGACGTGGCAGAAAC |
| CP | Fw-Asp | GAGGCAGAGTCACGTCTCAG |
| CP | Rv-Asp | TGAACGACACAGCGGATTTG |
| SQ | Fw-T3-Seq1-3 | GCAATTAACCCTCACTAAAGGCAAGCAGAAGACGGCATACGAGATCGTGATGTGACTGGAGTTCCTTGGCACCCGAGAATTCCAGGTCCTGCCCTTGACACATAC |
| SQ | Rv-T7-Seq2-2-*syy-1* | TAATACGACTCACTATAGGGCAAGCAGAAGACGGCATACGAGATACATCGGTGACTGGAGTTCCTTGGCACCCGAGAATTCCAACTGAGCGAAGGTCGAACTG |
| SQ | Rv-T7-Seq2-2-*syy-2* | TAATACGACTCACTATAGGGCAAGCAGAAGACGGCATACGAGATACATCGGTGACTGGAGTTCCTTGGCACCCGAGAATTCCACCACACCGAGGTTGTGACTTC |
| SQ | Fw-T3 | GCAATTAACCCTCACTAAAGG |
| SQ | Fw-T7 | TAATACGACTCACTATAGGG |
| NB | D-Ala | **Biotin-**ACCATCTGAGCTACATCCCC**-Biotin, HPLC-purified** |
| NB | D-Tyr | **Biotin-**TGCCGGAATCGAACCAGCG**-Biotin, HPLC-purified** |
| NB + WT | C-Tyr | **Biotin-**TACCGGATTCGAACCAGTG**-Biotin, HPLC-purified** |
| WS | C-Tyr | **Digoxigenin-**TACCGGATTCGAACCAGTG**-Digoxigenin, HPLC-purified** |
| SB | Fw-p | CTTGTACAGCTCGTCCATGC |
| SB | Rv-p | ATGGCCATCATCAAGGAGTT |

| **Aim(s)** | **Name** | **Description** |
|---|---|---|
| CL | GH1-sgRNA | Sequence of GH1 sgRNA. |
| CL | GH2-sgRNA | Sequence of GH2 sgRNA. |
| CL | Fw-pEF004-sgRNA-GH1 | Forward megaprimer to insert the GH1 sgRNA into the pEF004 vector by overlapping PCR (sgRNA-scaffold RNA). |
| CL | Rv-pEF004-sgRNA-GH1 | Reverse megaprimer to insert the GH1 sgRNA into the pEF004 vector by overlapping PCR (sgRNA-U6 promoter). |
| CL | Fw-pEF005-sgRNA-GH2 | Forward megaprimer to insert the GH2 sgRNA into the pEF005 vector by overlapping PCR (sgRNA-scaffold RNA). |
| CL | Rv-pEF005-sgRNA-GH2 | Reverse megaprimer to insert the GH2 sgRNA into the pEF005 vector by overlapping PCR (sgRNA-U6 promoter). |
| CL | LW1-sgRNA | Sequence of LW1 sgRNA. |
| CL | LW2-sgRNA | Sequence of LW2 sgRNA. |
| CL | LW3-sgRNA | Sequence of LW3 sgRNA. |
| CL | LW4-sgRNA | Sequence of LW4 sgRNA. |
| CL | LW5-sgRNA | Sequence of LW5 sgRNA. |
| CL | LW6-sgRNA | Sequence of LW6 sgRNA. |
| CL | Fw-atU3b-LW1-sgRNA | Forward primer to insert the LW1 sgRNA into the pAtU3b-sgR vector. |
| CL | Rv-atU3b-LW1-sgRNA | Reverse primer to insert the LW1 sgRNA into the pAtU3b-sgR vector. |
| CL | Fw-at7SL-LW2-sgRNA | Forward primer to insert the LW2 sgRNA into the pAt7SL-sgR vector. |
| CL | Rv-at7SL-LW2-sgRNA | Reverse primer to insert the LW2 sgRNA into the pAt7SL-sgR vector. |
| CL | Fw-atU6-LW3-sgRNA | Forward primer to insert the LW3 sgRNA into the pAtU6-sgR vector. |
| CL | Rv-atU6-LW3-sgRNA | Reverse primer to insert the LW3 sgRNA into the pAtU6-sgR vector. |
| CL | Fw-atU3b-LW4-sgRNA | Forward primer to insert the LW4 sgRNA into the pAtU3b-sgR vector. |
| CL | Rv-atU3b-LW4-sgRNA | Reverse primer to insert the LW4 sgRNA into the pAtU3b-sgR vector. |
| CL | Fw-at7SL-LW5-sgRNA | Forward primer to insert the LW5 sgRNA into the pAt7SL-sgR vector. |
| CL | Rv-at7SL-LW5-sgRNA | Reverse primer to insert the LW5 sgRNA into the pAt7SL-sgR vector. |
| CL | Fw-atU6-LW6-sgRNA | Forward primer to insert the LW6 sgRNA into the pAtU6-sgR vector. |
| CL | Rv-atU6-LW6-sgRNA | Reverse primer to insert the LW6 sgRNA into the pAtU6-sgR vector. |
| CL | Fw-pFK206-attB2-SV40NLS | Forward primer to amplify a recombinant pFK206 from attB2 site. Flanking sequence with SV40 NLS + STOP codon. |

| | | |
|---|---|---|
| CL | Rv-pFK206-attB1 | Reverse primer to amplify a recombinant pFK206 from attB1 site. |
| CL | Fw-attB1-pUBQ10-5'UTR | Forward primer to amplify Arabidopsis pUBQ10+5'UTR. Flanking sequence with attB1 site. |
| CL | Rv-pUBQ10-5'UTR | Reverse primer to amplify Arabidopsis pUBQ10+5'UTR. |
| CL | Fw-pUBQ10-5'UTR-EXT19CDS | Forward primer to amplify Arabidopsis EXT19 CDS. Flanking sequence with pUBQ10+5'UTR. |
| CL | Rv-EXT19CDS-2xGly | Reverse primer to amplify Arabidopsis EXT19 CDS. Flanking sequence with 2xGly linker. |
| CL | Fw-EXT19CDS-2xGly-GFPCDS | Forward primer to amplify GFP CDS. Flanking sequence with EXT19 CDS + 2xGly linker. |
| CL | Rv-GFPCDS-SV40NLS-STOP | Reverse primer to amplify GFP CDS. Flanking sequence with SV40 NLS + STOP codon. |
| CL | Fw-pUBQ10-5'UTR-mCherryCDS | Forward primer to amplify mCherry CDS. Flanking sequence with pUBQ10+5'UTR. |
| CL | Rv-mCherryCDS-SV40NLS-STOP | Reverse primer to amplify mCherry CDS. Flanking sequence with SV40 NLS. |
| CL | Fw-attB2-pFK206-p2S3-mCherry | Forward primer to amplify the pFK206-p2S3-mCherry vector with attB2 site and excluding the rbcs terminator. |
| CL | Fw-attB1-Ing-Y | Forward primer to amplify the engineered clustered $tRNA^{Tyr}$ gene strand with attB1 site. |
| CL | Rv-attB2-Ing-Y | Reverse primer to amplify the engineered clustered $tRNA^{Tyr}$ gene strand with attB2 site. |
| CL | Fw-attB1-1xSYY | Forward primer to amplify one repeat unit of the SYY cluster with attB1 site. |
| CL | Rv-attB2-1xSYY | Reverse primer to amplify one repeat unit of the SYY cluster with attB2 site. |
| GN | Fw-1 | Forward flanking primer to screen the presence/absence of the SYY cluster in *syy-1*/*syy-2* alleles. |
| GN | Rv-2-*syy-1* | Reverse flanking primer to screen the presence/absence of the SYY cluster in the *syy-1* allele. |
| GN | Rv-2-*syy-2* | Forward flanking primer to screen the presence/absence of the SYY cluster in the *syy-2* allele. |
| GN | Fw-3 | Forward flanking primer to screen the presence/absence of the SYY cluster in *syy-1*/*syy-2* alleles. |
| CP + GN | Fw-α | Forward internal primer to amplify the clustered $tRNA^{Ser}$ gene. |
| CP + GN | Rv-β | Reverse internal primer to amplify the clustered $tRNA^{Ser}$ gene. |
| CP + GN | Fw-γ | Forward internal primer to amplify the clustered $tRNA^{Tyr}$-1 gene. |
| CP + GN | Rv-δ | Reverse internal primer to amplify the clustered $tRNA^{Tyr}$-1 gene. |
| CP + GN | Fw-ε | Forward internal primer to amplify the clustered $tRNA^{Tyr}$-2 gene. |
| CP + GN | Rv-ζ | Reverse internal primer to amplify the clustered $tRNA^{Tyr}$-2 gene. |
| CP | Fw-Ala | Forward primer to amplify the control dispersed $tRNA^{Ala}$ gene. |
| CP | Rv-Ala | Reverse primer to amplify the control dispersed $tRNA^{Ala}$ gene. |
| CP | Fw-Val | Forward primer to amplify the control dispersed $tRNA^{Val}$ gene. |
| CP | Rv-Val | Reverse primer to amplify the control dispersed $tRNA^{Val}$ gene. |
| CP | Fw-Asp | Forward primer to amplify the control dispersed $tRNA^{Asp}$ gene. |
| CP | Rv-Asp | Reverse primer to amplify the control dispersed $tRNA^{Asp}$ gene. |
| SQ | Fw-T3-Seq1-3 | Forward primer to amplify the deletion zone of *syy-1* and *syy-2* alleles. |
| SQ | Rv-T7-Seq2-2-*syy-1* | Reverse primer to amplify the deletion zone of the *syy-1* allele. |
| SQ | Rv-T7-Seq2-2-*syy-2* | Reverse primer to amplify the deletion zone of the *syy-2* allele. |
| SQ | Fw-T3 | Forward T3 primer to sequence the deletion zone of *syy-1* and *syy-2* alleles. |
| SQ | Fw-T7 | Forward T7 primer to sequence the deletion zone of *syy-1* and *syy-2* alleles. |
| NB | D-Ala | Biotinylated probe reverse-complementary to dispersed $tRNAs^{Ala}$. |
| NB | D-Tyr | Biotinylated probe reverse-complementary to dispersed $tRNAs^{Tyr}$. |
| NB + WT | C-Tyr | Biotinylated probe reverse-complementary to clustered $tRNAs^{Tyr}$. |
| WS | C-Tyr | Digoxigenin-labelled probe reverse-complementary to clustered $tRNAs^{Tyr}$. |
| SB | Fw-p | Forward primer to amplify the p Southern blot probe. |
| SB | Rv-p | Reverse primer to amplify the p Southern blot probe. |

**CL:** cloning, **GN:** genotyping, **SQ:** sequencing, **CP:** chop-PCR, **NB:** northern blotting, **WT:** WISH-TSA, **WS:** WISH, **QP:** real-time quantitative PCR, **SB:** Southern blotting
